# The Reinstatement of a Forgotten Infantile Memory

**DOI:** 10.1101/2025.09.27.678956

**Authors:** Maria Lahr, Fabia Imhof, Lorenzo Mauro, Talia Ulmer, Flavio Donato

## Abstract

Infantile memories present a striking paradox: while early-life experiences are typically forgotten, reflecting the phenomenon of infantile amnesia, traumatic events from infancy can profoundly shape adult cognition and behavior. How do memories that are seemingly inaccessible persistently influence cognitive processes and behaviors throughout life? Rodent studies have demonstrated that forgotten infantile memories remain encoded as latent “infantile memory engrams” (iEngrams) within neuronal circuits, capable of memory reinstatement under artificial experimental conditions. Still, the network mechanisms underpinning the natural reinstatement of a forgotten infantile memory are unknown.

Here, we show that infantile memories, though physiologically irretrievable in adults, remain stored within hippocampal circuits and their engrams contribute to hippocampal network dynamics. Crucially, reinstating these memories requires a carefully orchestrated network process. An initial contextual reminder primes the hippocampal network to increase activity of the iEngram during a subsequent aversive reminder, which tags iEngram neurons for offline reactivation. This reactivation facilitates the integration of previously latent infantile memories with novel neuronal ensembles, reinstating behavior consistent with the original memory. These findings critically advance our understanding of the neuronal mechanisms underlying physiological memory encoding, retrieval, and reinstatement across development. Furthermore, they delineate the temporal boundaries and underlying physiological substrate of the process by which a latent representation becomes associated with a novel neuronal substrate, suggesting potential interventions to prevent the maladaptive reinstatement of traumatic infantile memories.

## Introduction

Infantile memories represent a paradox. On the one hand, memories encoded during development are typically inaccessible later in life due to infantile amnesia or the inability to consciously recall early-life events (*1–11*). On the other hand, early life events, especially traumatic ones, can exert profound and persistent effects later in life, shaping cognitive processes and emotional responses well into adulthood (*3, 6, 8, 12–15*). This observation raises a critical question: how can memories that appear forgotten continue to influence behavior throughout life? Rodent studies have provided key insights into this phenomenon, demonstrating that infantile memories are not erased during development but persist in a latent state, capable of modulating future behavior under appropriate conditions (such as those leading to memory reinstatement) (*1, 2, 4, 5, 7, 11, 16, 17*). Biological traces encoding these early-life experiences, termed infantile memory engrams (iEngrams), can evoke behavioral responses consistent with the original memory upon artificial activation, even though the physiological recollection of the memory fails when triggered by the relevant stimuli (*7, 17*). Yet, the physiological mechanisms enabling the natural reinstatement of a latent, forgotten infantile memory in the absence of artificial interventions remain unknown.

## Results

### The timely and sequential presentation of reminders reinstates a forgotten infantile memory in adulthood

To dissect the network mechanisms underlying the physiological reinstatement of a forgotten infantile memory, we adapted a contextual fear conditioning (cFC) protocol in infant mice (*11*). On postnatal day 19 (P19), animals were introduced to a novel environment (Training Context, TR) and exposed to a series of mild foot shocks (Fig. 1A, “cFC Training”). As previously shown, mice at this age can form associative memories for this event, as evidenced by elevated freezing upon re-exposure to the TR one day later (Recall 1d; Fig. S1A) (*1, 7, 9, 17*). However, these infant memories were transient and typically forgotten by adulthood (> P60), when mice no longer froze upon TR re-exposure (a hallmark of infantile amnesia. Recall > 40d, Fig. S1A).

**Fig. 1.**
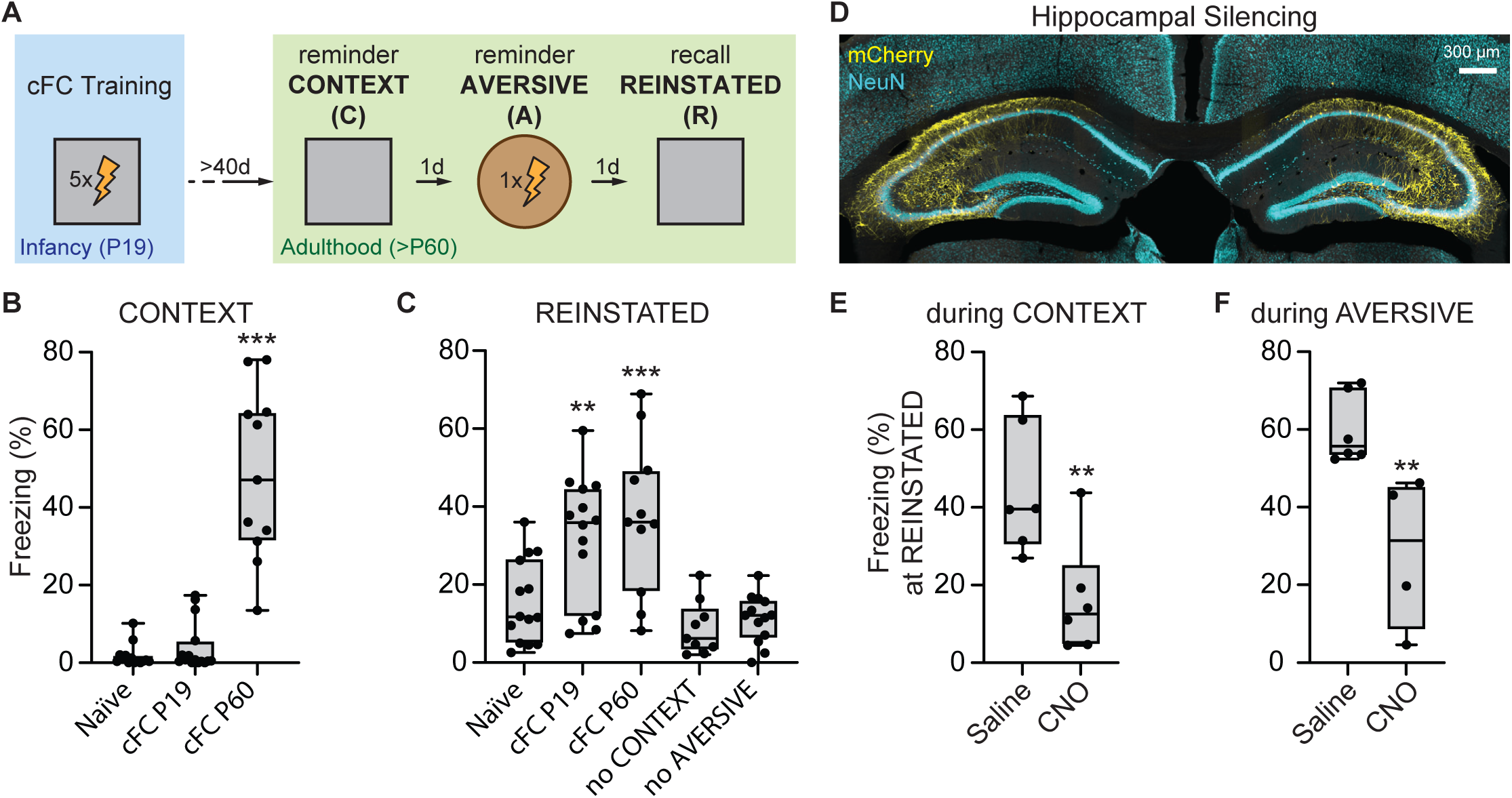
Hippocampus-dependent reinstatement of an infantile fear memory requires reminders of CONTEXT and AVERSIVE experiences. **(A)** Schematic of the behavioral protocol to trigger reinstatement of a forgotten infantile memory. Mice underwent contextual fear conditioning (cFC) training in infancy (P19), followed in adulthood by exposure to reminders of the CONTEXT and the AVERSIVE stimulus experienced during cFC training. The recall of the reinstated memory (REINSTATED recall) was tested in the CONTEXT one day after the final reminder. All behavioral sessions in adulthood were spaced by one day. **(B)** Freezing at CONTEXT reminder in adulthood was low in mice fear conditioned at P19, similar to animals that never experienced conditioning (Naϊve animals. Dunnett’s test, *p* = 0.634), and significantly higher in adult-conditioned animals (p < 0.001), indicating infantile amnesia. One-way ANOVA, F(2, 38) = 51.93, p < 0.001, r2 = 0.7321. n 2 11 mice per group. **(C)** At REINSTATED recall, mice conditioned at P19 expressed robust freezing (Dunnett’s test, p = 0.007), significantly higher than naϊve animals and comparable to animals conditioned during adulthood (> P60) only when both CONTEXT and AVERSIVE reminders were given. No reinstatement was observed in animals lacking either the CONTEXT (Dunnett’s test, p = 0.970) or the AVERSIVE (p > 0.999) reminder. One-way ANOVA, F(25, 226) = 11.18, p < 0.001, r2 = 0.5529; Brown-Forsythe test, F(25, 226) = 2.622, p < 0.001; Bartlett’s test, x2 = 81.17, p < 0.001. n 2 9 mice per group. **(D)** AAV-mediated expression of excitatory DREADD (hM3Dq-mCherry, yellow) in PV interneurons of dorsal hippocampus. Representative histological image showing mCherry expression in PV+ neurons in dorsal CA3 and CA1 (single-plane image, 10x objective; scale bar = 300 µm). **(E, F)** Activation of hippocampal PV interneurons via CNO injection during either the CONTEXT (E) or AVERSIVE (F) reminder prevented memory reinstatement, resulting in low freezing at REINSTATED recall. Student’s *t*-test against saline injected controls: CONTEXT, *t*(12) = 3.515, p = 0.004, Cohen’s d = 0.507; AVERSIVE, t(9) = 4.04, p = 0.003, Cohen’s d = 0.645. n = 6 for CONTEXT, n 2 4 for AVERSIVE.

To probe whether these latent memories could be reinstated physiologically, P19-trained mice remained undisturbed until adulthood (> P60), when they underwent a three-day reinstatement protocol (Fig. 1A). On day 1, mice were re-exposed to the TR without reinforcement (reminder “CONTEXT”, “C”). On day 2, they received a single foot shock in a distinct, novel environment (reminder “AVERSIVE”, “A”). On day 3, they were returned to the TR to assess whether the sequential reminders reinstated the expression of the infantile memory (recall “REINSTATED”, “R”).

As expected, P19-trained mice exhibited infantile amnesia: during the CONTEXT reminder, they froze at similarly low levels as naϊve controls (i.e., mice that had not learned cFC during infancy), and significantly less than P60-trained animals recalling an age-matched (remote) adult memory (Fig. 1B). Likewise, during the AVERSIVE reminder, their freezing response to the shock, and the strength of the newly formed memory for this event were indistinguishable from those of naϊve animals (Fig. S1B). Strikingly, during the REINSTATED recall session, P19-trained mice exhibited memory reinstatement and froze significantly more than naϊve animals, reaching levels comparable to P60-trained mice (Fig. 1C). Increased freezing during REINSTATED recall in P19-trained animals required the presentation of both reminders: omission of either the CONTEXT exposure or the foot shock presentation during the AVERSIVE reminder abolished reinstatement of the forgotten memory (Fig. 1C).

To identify which critical experiences triggered memory reinstatement, we tested the impact of each reminder and the temporal delay between their presentation. P19-trained mice failed to reinstate the memory if the CONTEXT reminder took place in a novel environment, both when memory testing at recall happened in the novel environment or the original training context, and even when partial sensory cues (e.g., the original training odor) were preserved in the novel environment (Fig. S2A). Likewise, replacing the foot shock in the AVERSIVE reminder with a distinct aversive experience (e.g., exposure to an elevated open platform) also failed to trigger reinstatement (Fig. S2A). Reversing the sequence of reminders, or presenting the same reminder twice without the other similarly disrupted memory reinstatement (Fig. S2A). Moreover, the infantile memory could be reinstated when the CONTEXT and AVERSIVE reminders were separated by up to three days, but not when the interval extended to seven or twenty-eight days (Fig. S2B). Last, reinstated freezing was observed when recall occurred 24 hours after the AVERSIVE reminder, but not after 6 or 15 hours (Fig. S2C). These findings suggest specific temporal constraints for reinstatement, necessitating both a maximum delay between reminders and a minimum consolidation period before expression.

Finally, we tested hippocampal contribution to memory reinstatement by silencing CA3 network activity during the presentation of the CONTEXT or AVERSIVE reminders using chemogenetic-mediated inhibition. AAVs encoding Cre-dependent hM3Dq DREADDs were injected into the hippocampus of PV-Cre mice, enabling selective activation of parvalbumin-positive interneurons and suppression of local excitatory activity (Fig. 1D). Administration of CNO 40 minutes before either the CONTEXT or AVERSIVE reminders impaired memory reinstatement, as shown by significantly reduced freezing during REINSTATED recall compared to controls (Fig. 1E and F, respectively). Together, these results demonstrate that the reinstatement of a forgotten infantile memory is a hippocampus-dependent process triggered by the temporally constrained, sequential presentation of specific reminders of the original context and aversive stimulus experienced during infancy.

### Hippocampal population dynamics underpinning forgotten memory reinstatement

We next investigated neuronal recruitment and population dynamics underlying the reinstatement of the forgotten infantile memory, focusing on the contribution of infantile engram (iEngram) neurons. To label iEngram cells, we exploited a transgenic mouse line expressing CreERT2 recombinase driven by the c-Fos promoter (TRAP-tdTomato) (*18*). Administration of 4-hydroxytamoxifen (4-OHT) immediately after cFC training at P19 permanently tagged neurons activated during memory encoding with the fluorescent marker tdTomato (TRAPed cells, Fig. S3A and B). In adult mice, we mapped iEngram neurons recruitment to c-Fos+ ensembles by quantifying the overlap between tdTomato and c-Fos expression at key phases of the reinstatement protocol. Specifically, we assessed c-Fos protein expression in iEngram cells after the CONTEXT and AVERSIVE reminders, as well as during the following Early (3-5 h), Mid (7-9 h), and Late (13-15 h) offline periods that mice spent in the home cage (Fig. S3A). Broad hippocampal engagement, reflected by increased c-Fos expression in DG, CA3, and CA1, was observed following either reminder and during corresponding Late offline periods (C-Late and A-Late, respectively Fig. S3C-E, left), consistent with known consolidation windows (*19–21*). Notably, this network-wide activation pattern occurred even upon presentations of reminders that failed to produce memory reinstatement (Fig. S3C-E, center-left). In contrast, selective c-Fos expression in iEngram neurons was observed specifically in the DG and CA3 areas of the hippocampus, and exclusively under conditions that led to successful memory reinstatement (Fig. S3C-E, right). Specifically, iEngram recruitment to c-Fos+ ensembles in DG and CA3 was higher than expected by chance upon the CONTEXT reminder and the Late offline period following the AVERSIVE reminder (Fig. S3C-E, center-right), but at chance level when only one reminder was presented (Fig. S3C-E, right). Together, these results reveal that while the hippocampal network responds to individual reminders of infantile experience with a broad network engagement, the reactivation of iEngram neurons follows more restricted recruitment, with selective c-Fos expression upon behavioral protocols leading to successful memory reinstatement.

While increased c-Fos expression suggests that the reactivation of iEngram cells in DG and CA3 is a key neural correlate of successful reinstatement of a forgotten infantile memory, they fall short of revealing whether activity dynamics in individual iEngram neurons follow any specific longitudinal patterns spanning the presentation of the reminders and the offline consolidation windows necessary for successful reinstatement. To address this specific question, we used calcium imaging to track individual hippocampal CA3 neurons across the reinstatement protocol (*21*). We focused on CA3 due to its prominent c-Fos expression in iEngram neurons (Fig. S3D), as well as its established role in memory retrieval. We combined TRAP labeling to tag iEngram neurons with a virally-expressed GCaMP6f to monitor calcium activity in both iEngram (tdTomato+/GCaMP6f+) and non-iEngram CA3 cells (tdTomato-/GCaMP6f+) across the same timepoints delineated previously (Fig. 2A-C). At the start of each recording day, we acquired a brief baseline session in the home cage to normalize activity levels across days and account for drifting properties of individual neurons (Fig. 2A). We then computed an “Activation Score” (AS) to classify each neuron as activated, neutral or inhibited by comparing its average activity rage across the whole recording period during a behavioral or offline session, relative to its average baseline recording on each day (*21, 22*). Neurons with sustained activation during the CONTEXT or AVERSIVE reminders were assigned to their respective ensembles (Fig. 2D), enabling comparisons of their dynamics with the broader CA3 network and iEngram cells.

**Fig. 2.**
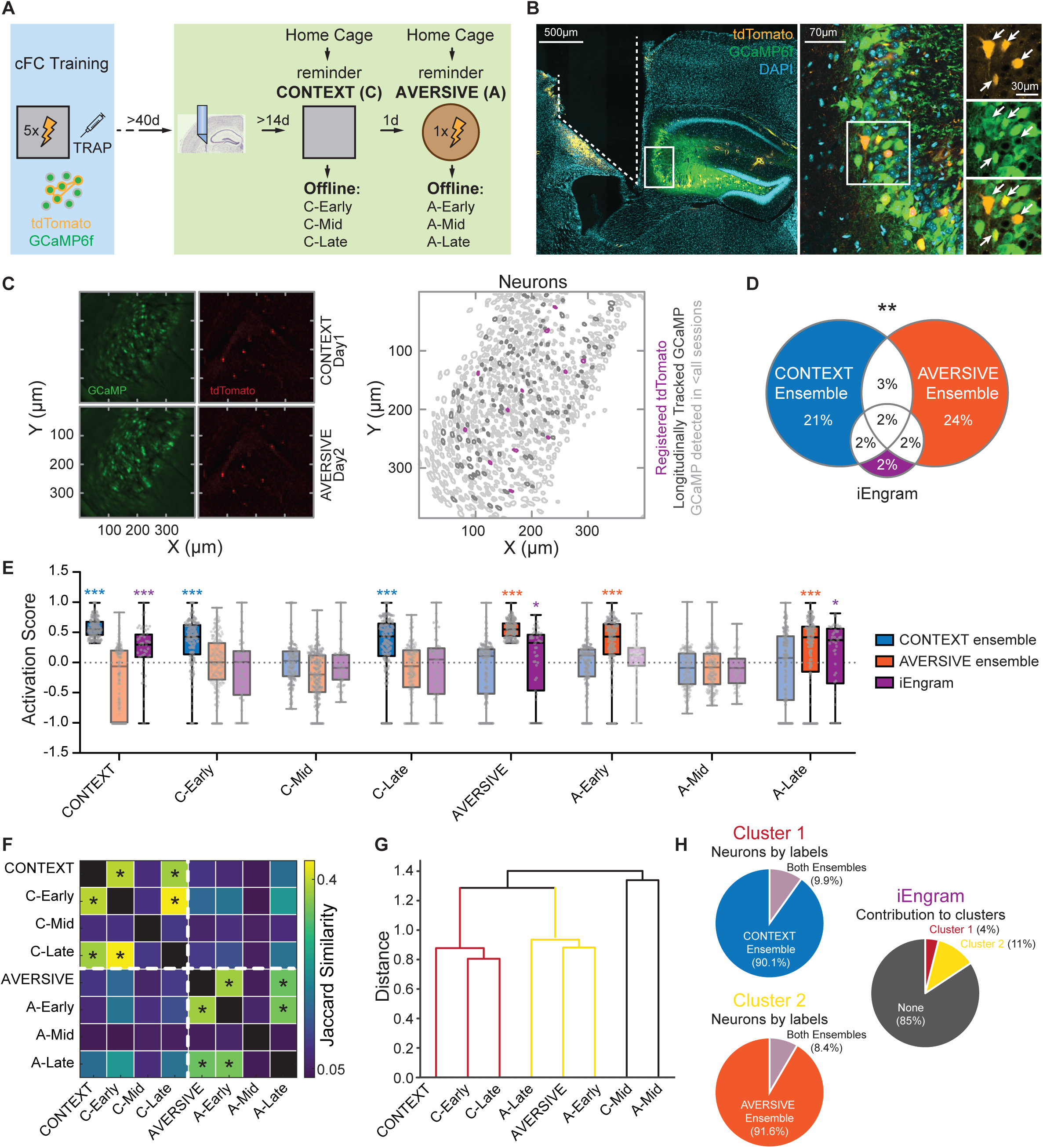
CA3 network activity during forgotten infantile memory reinstatement. **(A)** Schematic of the experimental design. iEngram neurons were permanently labeled via 4-OHT-induced TRAPing of Fos+ cells during infant cFC Training (P19) in TRAP2 mice, where an AAV vector for the pan-neuronal expression of the calcium indicator GCaMP6f mixed with a Cre dependent tdTomato vector, had been previously locally infused into hippocampal CA3 on P2. In adulthood, mice underwent prism implantation adjacent to CA3, to perform dual-color, one-photon calcium imaging during CONTEXT and AVERSIVE reminders and offline sessions. Baseline recordings were performed daily in the home cage before behavior. Offline activity was assessed at +3-5 h (Early), +7-9 h (Mid), and +13-15 h (Late) following each behavioral session. **(B)** Representative histology showing the implantation site of the microendoscope (DAPI, cyan), expression of tdTomato in iEngram cells (yellow), and pan-neuronal expression of GCaMP6f (green) in the CA3 pyramidal layer. Single-plane image (10x objective, scale bar scale bar = 500 µm); magnified single-plane images (30x objective, scale bar = 70 µm) and zoom in images (scale bar = 30 µm) show co-labeled iEngram neurons (white arrows). **(C)** Left: Maximum intensity projections of calcium signals acquired during CONTEXT (upper row, left) and AVERSIVE (lower row, left) reminders, and recorded tdTomato-labeled iEngram neurons (right). Right: Individual neurons were longitudinally tracked across all sessions. Regions of interest (ROIs) were color-coded: iEngram neurons (magenta), neurons tracked across all sessions (dark grey), and those tracked in a minority of sessions (light grey). **(D)** Neural ensembles were defined by activation scores (AS) during CONTEXT or AVERSIVE reminders, or by the expression of tdTomato (iEngram). Despite including similar proportions of all longitudinally tracked neurons (21 and 24%, respectively), CONTEXT and AVERSIVE ensembles were orthogonalized on largely non-overlapping neuronal subsets (Monte Carlo simulations, p = 0.002, Cohen’s d = −2.88) with only 3% of all tracked neurons belonging to both ensembles. iEngram neurons equally contributed to each category. **(E)** Distinct activation profiles across the reinstatement protocol for CONTEXT, AVERSIVE, and iEngram neurons. CONTEXT ensembles were activated during CONTEXT reminder and its Early and Late offline phases, while AVERSIVE ensembles were activated similarly around AVERSIVE reminder and its Early and Late offline phases. iEngram neurons responded with increased activity to both reminders and were selectively active in the Late offline phase following AVERSIVE. n = 690 neurons from 7 mice; repeated-measures ANOVA with Geisser-Greenhouse correction: CONTEXT, F(5.105, 934.2) = 69.81, p < 0.001, 12 (partial) = 0.276, £ = 0.7293; AVERSIVE, F(5.011, 972.1) = 71.80, p < 0.001, 12 (partial)= 0.270, £ = 0.7158. iEngram neurons, F (5.345, 261.9) = 2.909, p = 0.012, 12 (partial) = 0.215, £ = 0.7636. Significance reported according to a two-tailed, one-sample Wilcoxon signed-rank test relative to baseline activation. Recordings where the two-tailed one-sample Wilcoxon Signed-Rank test revealed no significant activation compared to baseline are transparent. **(F)** Jaccard similarity of active populations across sessions revealed session pairs where the fraction of neurons that were activated during both sessions was higher than expected by chance based on a permutation test (significance beyond the 95^th^ percentile of the null distribution based on shuffled matrices, white lines highlight similarity clustering). **(G)** Hierarchical clustering of Jaccard distances revealed two major activity clusters corresponding to CONTEXT (Cluster 1) and AVERSIVE (Cluster 2) experiences and their associated Early and Late offline periods. **(H)** Matching neurons’ tuning classification to cluster membership revealed a strong association between functional identity and ensemble structure. Specifically, 90.1% of neurons in Cluster 1 belonged to the CONTEXT ensemble, while 91.6% of neurons in Cluster 2 were part of the AVERSIVE ensemble. iEngrams contributed modestly to both clusters (Cluster 1: 4%, Cluster 2: 11%). These associations were statistically robust for both clusters (Cluster 1: x2(3) = 243.76, p < 0.001, Cramer’s V = 0.944; Cluster 2: x2(3) = 194.37, p < 0.001, Cramer’s V = 0.883), indicating a near-perfect correspondence between cluster identity and activation category.

CA3 network activity broadly mirrored c-Fos results, with elevated activity during reminder sessions and during Early and Late offline periods (Fig. S4A). CONTEXT and AVERSIVE ensemble neurons showed more restricted activation profiles: elevated activity during their corresponding reminder and its associated offline periods, but baseline or suppressed activity at other times (Fig. 2E and S4B-C). In contrast, iEngram neurons activation was elevated during both reminder sessions; during offline periods, activation was restricted to the Late recording following the AVERSIVE reminder (Fig. 2E and S4D). Moreover, although the CONTEXT and AVERSIVE ensembles were largely composed by non-overlapping sets of neurons (Fig. 2D), iEngram neurons were distributed across both, with subsets recruited to either one, both, or neither. This pattern suggests that iEngram neurons represent a diverse population engaged at multiple phases of the reinstatement protocol.

To probe neuronal co-activation patterns across sessions at the single-cell level, we calculated session-to-session overlap in activated neurons. Observed co-activity exceeded chance levels for specific session pairs (Fig. S5 A and B). Jaccard similarity among active population vectors calculated over entire sessions (Fig. 2F) revealed two major session clusters: one centered on the CONTEXT reminder day and its offline home-cage periods (Cluster 1), and the other centered on the AVERSIVE reminder day (Cluster 2) (Fig. 2G). Mean within-cluster similarity values (0.40 and 0.36) were significantly higher than expected by chance (p < 0.001, permutation test, Fig. S5C). We next identified neurons active across all conditions within each cluster. CONTEXT ensemble neurons were overrepresented in Cluster 1 (x2 p < 1e-38) and AVERSIVE ensemble neurons dominated Cluster 2 (x2 p < 1e-31), with minimal overlap, indicating that CA3 encodes the two reminders in orthogonal, time-locked ensembles (Fig. 2H, left). Longitudinal tracking confirmed that individual CONTEXT or AVERSIVE neurons preferentially contributed to their respective clusters (Fig. S4 A and B, right). By contrast, the majority of iEngram neurons exhibited cross-cluster activity, with stronger involvement in the AVERSIVE-associated cluster (Fig. 2H, right and S4D), suggesting they may serve to bridge temporally and functionally distinct memory components.

### iEngram activity during aversive reminder presentation and offline consolidation supports memory reinstatement

The activation of CA3 iEngram neurons by both CONTEXT and AVERSIVE reminders prompted us to test whether their activity was required to reinstate the forgotten infantile memory. To address this, we performed a series of loss-of-function experiments targeting CA3 iEngram neurons using the TRAP technology. First, we induced the expression of Cre-dependent inhibitory effectors, either the optogenetic silencer ArchT or the chemogenetic receptor hM4Di, by administering 4-OHT immediately after cFC training at P19. In adulthood, we selectively suppressed iEngram activity during specific phases of the reinstatement protocol, using either light delivery via implanted optic fibers (ArchT-expressing animals) or systemic CNO administration (hM4Di-expressing animals).

We first used optogenetics to restrict silencing to the CONTEXT or AVERSIVE reminder sessions and assessed reinstatement by measuring freezing during the REINSTATED recall session (Fig. 3A and B). Silencing iEngram neurons during the AVERSIVE reminder - but not during the CONTEXT reminder - prevented memory reinstatement (Fig. 3C and D). Only in the AVERSIVE-silenced group the strength of reinstatement across individual animals was inversely correlated with the proportion of ArchT-expressing iEngram neurons in CA3 (Fig. S6C), suggesting that iEngram activity during the AVERSIVE reminder is critical for memory reinstatement. This effect was specific to iEngram neurons TRAPed at P19 during memory encoding: silencing a similarly sized, nonspecific CA3 population labeled based on c-Fos expression during unrelated home cage activity in infancy had no impact on reinstatement in adults (Fig. 3D and S6D).

**Fig. 3.**
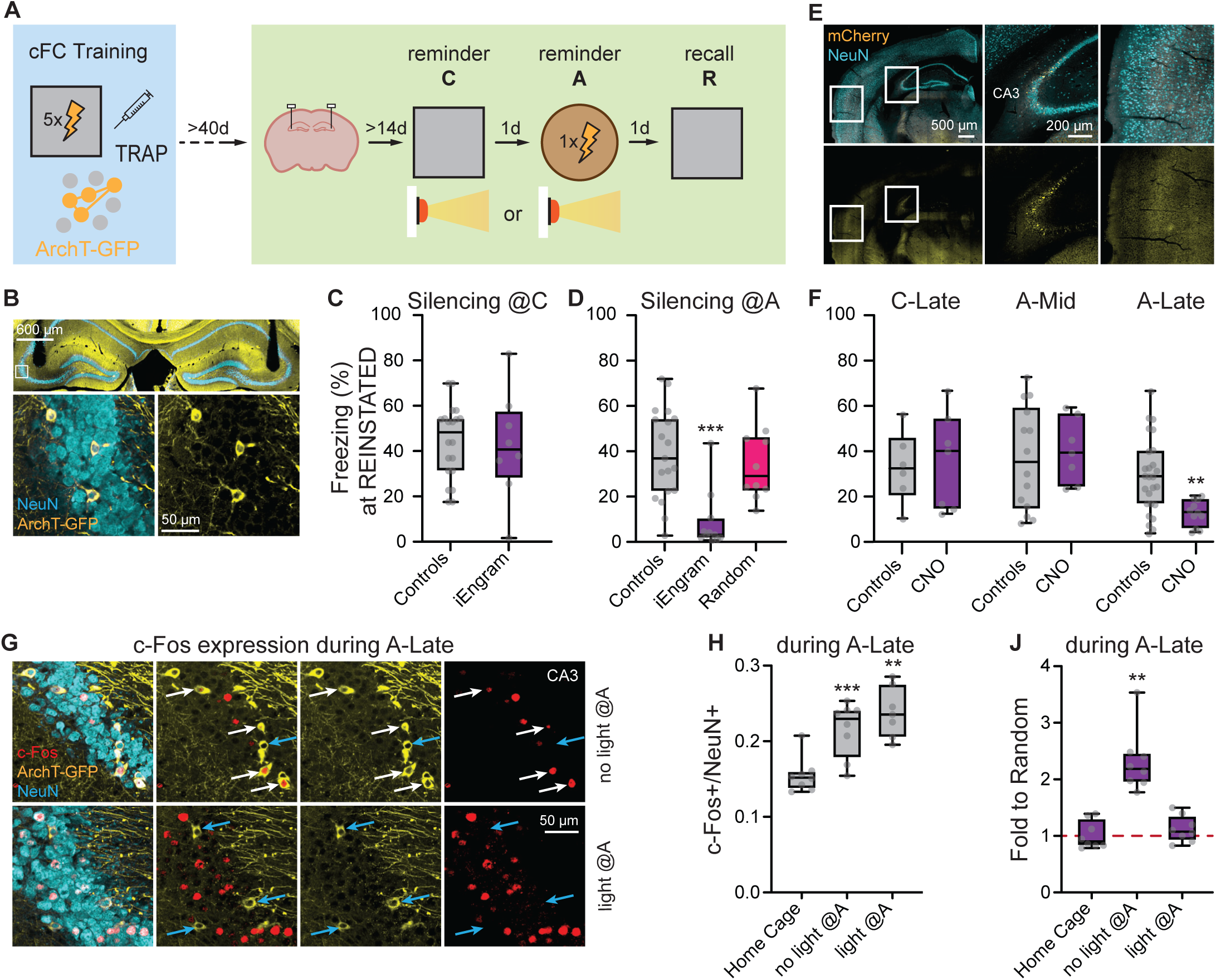
Activation of iEngram neurons during AVERSTVE reminder and associated Late offline period is necessary for infantile memory reinstatement. **(A)** Schematic of the experimental design. During cFC in infancy, iEngram neurons were genetically labeled (TRAPed) to express ArchT-GFP. In adulthood, optic fibers were bilaterally implanted above CA3 to allow light delivery during the CONTEXT or AVERSIVE reminder sessions for optogenetic inhibition. **(B)** Representative histology showing optical fiber tracks (upper panel, single-plane image, 10x objective, scale bar = 600 µm), and expression of ArchT-GFP in CA3 neurons (lower panel, single-plane images, 30x objective, Scale bar = 50 µm). **(C)** Light-mediated silencing of iEngrams during the CONTEXT reminder did not alter freezing levels at REINSTATED recall compared to controls (a mix of non-implanted mice, and animals expressing ArchT-GFP in iEngram neurons in the absence of light delivery during reminders. Two-tailed Mann-Whitney test, U = 74, p = 0.7745, r = 0.058, n 2’ 8 per group). **(D)** Silencing of iEngram neurons during the AVERSIVE reminder session prevented memory reinstatement, with reduced freezing in iEngram-TRAPed animals compared to animals where a random ensemble was TRAPed while mice were in their home cage days after cFC training during infancy. Control group as in (C). (Kruskal-Wallis test, H(2) = 15.60, p < 0.001, 12 = 0.368; Dunn’s test: iEngram vs. Controls, p < 0.001; Random vs. Controls, p = 0.303). n 2’ 10 per group. **(E)** Histological verification of hM4Di-mCherry expression in hippocampal CA3 (Single-plane image, 10x objective, scale bar = 500 µm). Overview image shows mCherry expression (yellow) over pan-neuronal staining (NeuN, cyan). Expression was confined to TRAPed neurons in the hippocampal pyramidal layer (middle panel), with no labeled somata in other regions (background likely due to axonal processes, right panel, scale bar = 200 µm). **(F)** Behavioral analysis revealed that chemogenetic inhibition of iEngrams during C-Late (n = 7) and A-Mid (n = 7) offline periods did not affect freezing at REINSTATED recall compared to controls (two-tailed Mann-Whitney tests: C-Late: U = 19, p = 0.836, r = 0.079; A-Mid: U = 45, p = 0.799, r = 0.065). In contrast, inhibition during A-Late (n 2’ 10) significantly reduced freezing compared to controls (U = 53, p = 0.002, r = 0.492). Controls were a mixture of CNO-injected animals; DREADD-expressing, saline-injected animals; and DREADD-negative, not-injected animals (n = 47). **(G)** Representative histology showing ArchT-GFP TRAPed iEngram neurons (yellow) and c-Fos (red) expression during A-Late offline periods in controls (no light during AVERSIVE, top row) or experimental (light during AVERSIVE, bottom row) animals. White arrows indicate GFP+/c-Fos+ neurons, while blue arrows indicate GFP+/c-Fos-neurons. Notice the higher prevalence of double-positive neurons in control conditions than upon silencing of iEngram during AVERSIVE reminder (Single-plane images, 30x Objective, Scale bar = 50 µm). **(H, J)** Inhibition of iEngram neurons during the AVERSIVE reminder did not affect the overall increase in c-Fos+ neuron prevalence in the CA3 network during the following A-Late offline phase (one-way ANOVA, F(2, 20) = 18.67, p < 0.001, 12 = 0.651; Dunn’s test: No Light vs. Home Cage, p < 0.001; Light vs. Home Cage, p = 0.002;) (H). However, this manipulation significantly prevented c-Fos expression specifically in iEngram neurons at A-Late (one-way ANOVA, F(2, 21) = 29.75, p < 0.001, 12 = 0.7391; Dunn’s test: No Light vs. Home Cage, p < 0.001; Light vs. Home Cage, p = 0.792. Two-tailed one-sample Wilcoxon Signed Rank Test revealed no change in c-Fos expression in TRAPed neurons upon optogenetic silencing, in contrast to increased expression observed in no light controls) (J).

We next tested whether iEngram activity during the Late offline period following the AVERSIVE reminder was also necessary for memory reinstatement. To selectively silence these neurons during this extended window, we used chemogenetics (Fig. 3E and S7A). Mice expressing hM4Di in CA3 iEngram neurons received CNO to induce peak silencing during the Late offline phase after the AVERSIVE reminder. As controls, we silenced iEngram neurons during periods not associated with elevated activity or c-Fos expression: specifically, the Late offline period after the CONTEXT reminder and the Mid offline period after the AVERSIVE reminder. Silencing during the Late offline period after the AVERSIVE reminder selectively impaired memory reinstatement, while silencing at other times had no significant effect on behavior (Fig. 3F).

These findings showed that successful reinstatement relied on iEngram activation during both the AVERSIVE reminder and in the subsequent Late offline period. This suggested that these two phases may be functionally connected: specifically, that iEngram activity during the reminder could “tag” the engram for later recruitment during offline consolidation. To test this, we optogenetically silenced iEngram neurons during the AVERSIVE reminder and assessed their recruitment to c-Fos+ ensembles during the Late offline period (Fig. S7B). While the overall increase of c-Fos expression in the broader CA3 network upon A-Late was unaffected by iEngram silencing during AVERSIVE reminder (Fig. 3H), c-Fos expression in iEngram neurons was prevented and at chance levels (Fig. 3J), suggesting that iEngram activity during the AVERSIVE is necessary to tag the infantile CA3 memory trace for later incorporation into offline memory ensembles during consolidation.

### The expression of the forgotten infantile memory is supported by a distinct CA3 neuronal ensemble

Prompted by the observation that reinstated memories exhibited adult-like properties such as long-term persistence and specificity (unlike the transient, generalized infant memory, Fig. S8A and B), we hypothesized that reinstatement may trigger the formation of a new neuronal ensemble supporting recall. To test this, we analyzed c-Fos expression and calcium activity in CA3 during the REINSTATED session (Fig. 1A).

CA3 network activation was significantly elevated during REINSTATED recall, as reflected by both increased c-Fos+ neurons prevalence (Fig. 4A, left) and higher population-level activity rates (Fig. S8C). However, iEngram neurons showed no such increase in c-Fos expression (Fig. 4A, right) or activity (Fig. S8C) pointing out they are not part of activated ensembles during this session. This was further supported by optogenetic silencing experiments: inhibiting iEngram neurons during REINSTATED recall did not impair memory retrieval (Figure 4B). Together, these results strongly support the idea that the expression of the reinstated memory does not rely on reactivation of the original infant engram but rather on distinct neuronal ensembles, possibly emerging during the reinstatement protocol. We thus identified a “REINSTATED” ensemble, comprising neurons that were not part of the CONTEXT or AVERSIVE ensembles (Fig. 4C and D), and exhibiting selective activation during the REINSTATED session and the Late offline period following the AVERSIVE reminder (Fig. 4D and S9A, B). Co-activity analyses confirmed that the REINSTATED session did not cluster with prior ensemble dynamics (Clusters 1 or 2), and it exhibited the highest similarity with the Late offline session after the AVERSIVE reminder (Fig. 4E, left and center, and S10A and B). This similarity was driven by activation of REINSTATED ensemble neurons (Fig. 4E right). To test whether this new ensemble was causally involved in the expression of the reinstated memory, we TRAPed neurons active during the Late offline period after the AVERSIVE reminder using 4-OHT, thereby inducing ArchT expression specifically in this population (Fig. 4F). c-Fos analysis revealed that A-Late TRAPed neurons were selectively recruited to c-Fos+ ensembles during REINSTATED recall performed 7 days post AVERSIVE, while c-Fos+ expression in iEngram neurons remained at chance levels as previously reported for shorter delays (Fig. 4G and A, respectively). Optogenetic silencing of A-Late TRAPed neurons during the REINSTATED session impaired memory recall in proportion to the fraction of labeled cells (Fig. 4H, S11A-C), whereas silencing a comparable fraction of iEngram neurons, or neurons active during earlier offline windows, had no effect.

**Fig. 4.**
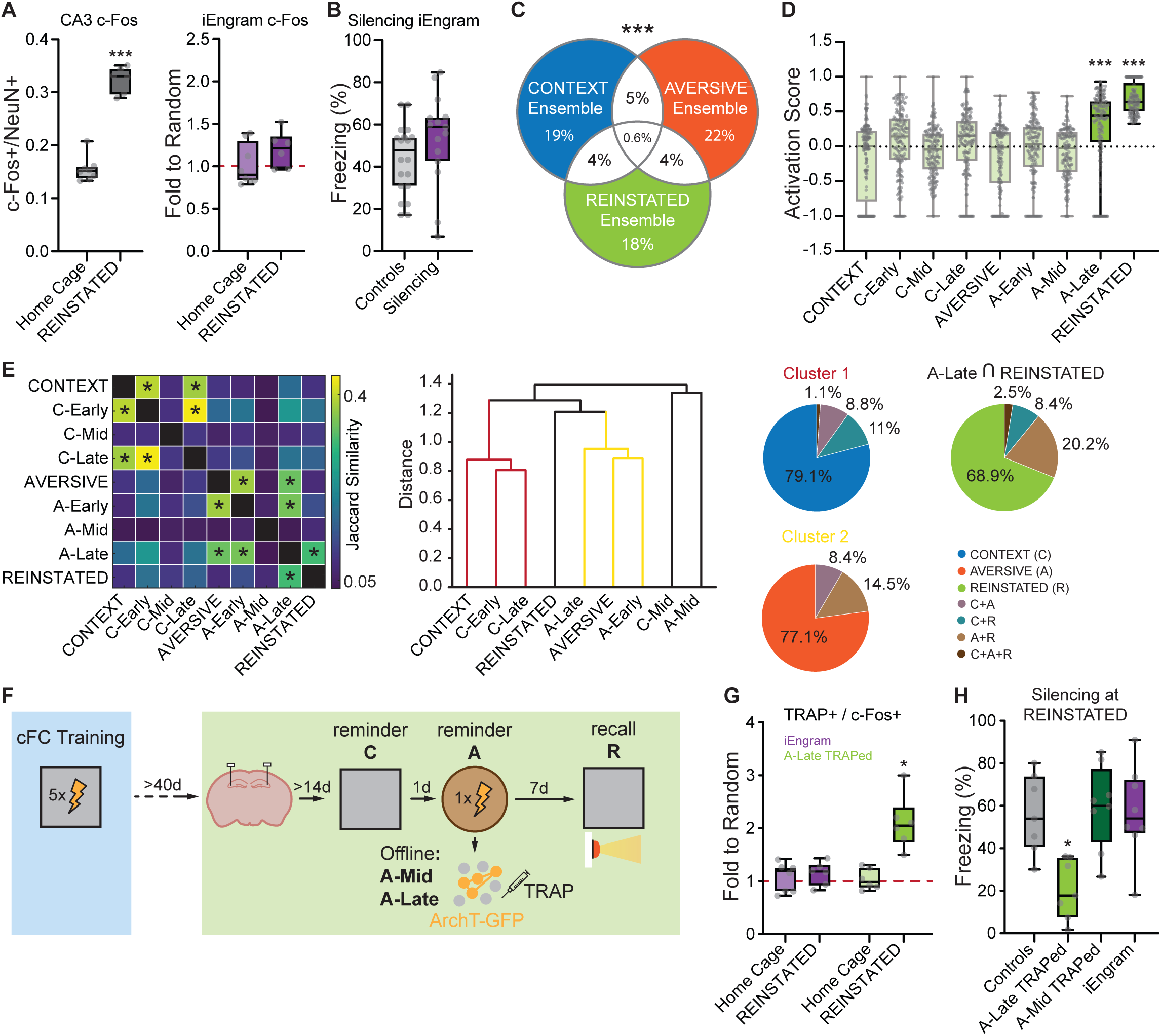
Representation of the REINSTATED memory by a distinct newly formed neuronal ensemble. **(A)** c-Fos expression in the CA3 network was significantly increased during REINSTATED recall (left), but expression in iEngram neurons was not elevated compared to chance (right) (Mann-Whitney tests: total c-Fos, U = 0, p < 0.001, r = 0.828; iEngram c-Fos, U = 14, p = 0.228, r = 0.345. Two-tailed one-sample Wilcoxon Signed Rank Test revealed no change in c-Fos expression in TRAPed neurons to what expected by chance. n 2: 6 per group). **(B)** Optogenetic inhibition of iEngram neurons during REINSTATED recall did not prevent the expression of the reinstated memory (controls is a pool of non-implanted animals, and ArchT-expressing animals where no light was delivered at recall. Two-tailed Mann-Whitney test, U = 96, p = 0.0727, r = 0.304; n 2: 15 per group). **(C)** Neurons with increased activity rates (AS > 0.33) during REINSTATED recall were defined as the REINSTATED ensemble. The REINSTATED ensemble displayed a unique activation profile: selective activation during REINSTATED recall and the Late offline phase following the AVERSIVE reminder. Repeated-measures ANOVA with Geisser-Greenhouse correction: F(6.129, 1011) = 58.26, p < 0.001, 12 = 0.261, £ = 0.7661. Significance reported according to a two-tailed, one-sample Wilcoxon signed-rank test relative to baseline activation. Recordings where the two-tailed one-sample Wilcoxon Signed-Rank test revealed no significant activation compared to baseline are transparent. **(D)** Despite including similar proportions of all longitudinally tracked neurons to the CONTEXT and AVERSIVE ensembles, the REINSTATED ensemble was orthogonalized onto a yet distinct subset of neurons (Monte Carlo simulations, p < 0.001 for each comparison). **(E)** Left: Jaccard similarity analysis revealed low similarity between activity patterns during REINSTATED recall and prior sessions, except for A-Late. Center: A separate lineage emerged for REINSTATED recall, distinct yet closest to Cluster 2, with the greatest similarity to A-Late. Right: the similarity between REINSTATED and A-Late was driven primarily by neurons identified as REINSTATED ensemble members (Monte Carlo simulation: Z = −4.54, p < 0.001, Cohen’s d = −4.54). **(F)** Schematic of the protocol for TRAP-based labeling of neurons active during the A-Late phase and their optogenetic inhibition during REINSTATED recall. Notice the 7-day delay between the AVERSIVE reminder and REINSTATED recall, to allow for expression of the TRAP-induced payload (ArchT). **(G)** c-Fos expression was significantly elevated in the A-Late TRAPed neurons during REINSTATED recall (Mann-Whitney comparison to Home Cage levels: U = 0, p = 0.002, r = 0.831. Two-tailed one-sample Wilcoxon Signed Rank Test revealed significant increase in c-Fos expression in A-Late TRAPed neurons upon REINSTATED recall, W = 21, p = 0.031), whereas iEngram neurons showed c-Fos expression at chance levels, consistent with Home Cage controls (Mann-Whitney comparison to Home Cage levels: U = 14, p = 0.366, r = −0.277. Two-tailed one-sample Wilcoxon Signed Rank Test W = 13, p = 0.218). n 2: 6 mice. **(H)** Optogenetic inhibition of TRAPed A-Late neurons during REINSTATED recall led to reduced freezing compared to controls (controls as in Fig. 3B, Kruskal-Wallis test, H(3) = 12.40, p = 0.006, 12 = 0.362). Post hoc Dunn’s tests confirmed a significant reduction for A-Late inhibition (p = 0.023), whereas inhibition of neurons TRAPed during A-Mid (p > 0.999) or iEngram neurons (p > 0.999) did not affect freezing. n 2: 7 mice.

Finally, we tested whether behavioral interference during the critical offline A-Late window could disrupt REINSTATED ensemble formation and prevent reinstatement. Mice underwent a protocol mimicking extinction, whereby they were exposed to the original training context for 30 minutes beginning at about 15 hours after the AVERSIVE reminder (Fig. S12A). This experience abolished reinstatement, with freezing levels comparable to naϊve controls (Fig. S12B). The effect was context-specific since exposure to a different context had no effect on reinstatement (Fig. S12B). Baseline levels of freezing upon A-Late behavioral interventions were still observed 7 days following the AVERSIVE reminder, ruling out the possibility of delayed reinstatement (Fig. S12B). Thus, the Late post-AVERSIVE offline window emerges as the critical moment when new neuronal ensembles are bound to a latent representation for the reinstatement of a forgotten infant memory, and as a possible critical window for behavioral interventions aimed at preventing reinstatement.

### Co-activity of iEngram and REINSTATED ensembles during Synchronous Network Events correlates with effective memory reinstatement

Having established that iEngram reactivation during the Late offline period after the AVERSIVE reminder (A-Late) is critical for memory reinstatement, and that neurons TRAPed during this window become incorporated into the reinstated memory trace, we next investigated how A-Late activity dynamics drive the formation of this new ensemble. Previous studies show that synchronous calcium events (SCEs), brief bursts of coordinated network activity (*23, 24*) (Fig. 5A), can link distinct memory traces. We therefore hypothesized that repeated co-activation of iEngram neurons with a separate neuronal population during A-Late SCEs could similarly tether this new ensemble to the latent infantile memory.

**Fig. 5.**
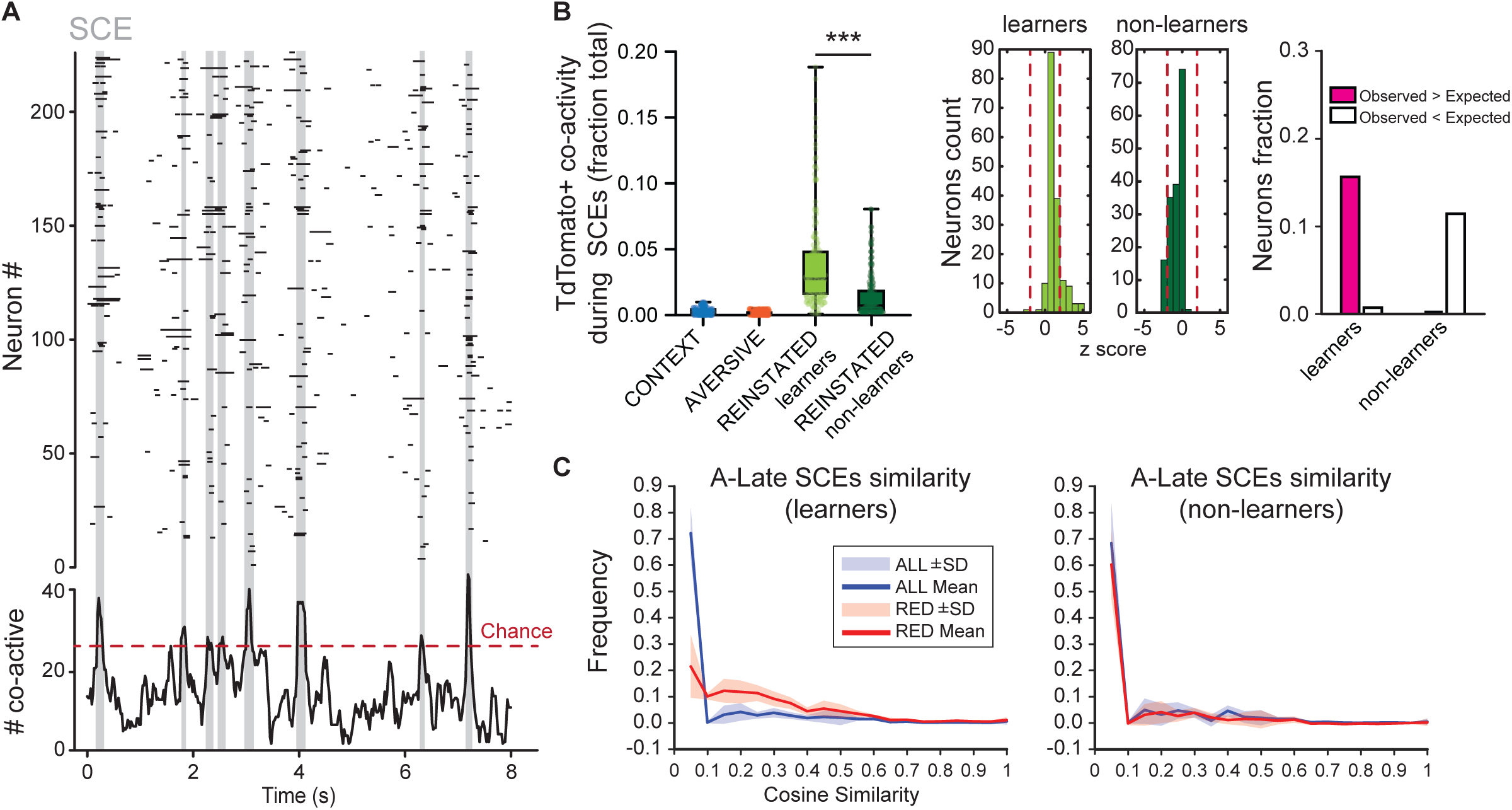
Co-activity of iEngram and REINSTATED ensembles during network events. **(A)** Top: Raster plot of neuronal activity during a representative offline session. Each row represents a neuron; ticks indicate active frames. Shaded gray areas mark synchronous calcium events (SCEs), defined as frames where the number of active neurons exceeded chance. Bottom: Line diagram showing the number of active neurons per frame. The dashed line indicates the session-specific significance threshold calculated on shuffle-based null models. Values above this threshold correspond to synchronous events shown in the raster plot. **(B)** Left: Box-and-whisker plot showing, for each ensemble (CONTEXT, AVERSIVE, REINSTATED learners, REINSTATED non-learners), the fraction of red SCEs (i.e., events with at least one tdTomato+ neuron active) in which each neuron was active during the A-Late period. Each dot represents one neuron. Statistical comparisons (Kruskal-Wallis: H(3) = 317.1, p < 0.001, £2 = 0.44) revealed significantly higher recruitment in REINSTATED learners compared to CONTEXT (Dunn’s multiple comparisons test, p < 0.001) and AVERSIVE (p < 0.001) ensembles. REINSTATED non-learners were also significantly different from CONTEXT (p = 0.008) and AVERSIVE (p < 0.001); CONTEXT vs AVERSIVE was not significant (p = 0.168). Notably, neurons in the REINSTATED learner ensemble were active in a significantly greater fraction of red SCEs compared to non-learners (p < 0.001). Center: distribution of z-scores for neuronal participation in red SCEs, computed per condition using a shuffle-based null model. Red dashed lines indicate the ±1.96 threshold (corresponding to p < 0.05, two-tailed). X-axis ranges from −5 to +5 to facilitate comparison. Right: grouped bar plots showing the proportion of neurons significantly above (pink) or below (white) expectation (p < 0.05) for each reinstatement condition. REINSTATED learners had more neurons participating above (z = 6.81, p < 0.001, h = 0.36) and fewer below (z = −2.81, p = 0.005, h = −0.30) expectation. In contrast, REINSTATED non-learners had fewer above (z = −3.20, p = 0.001, h = −0.45) and more below (z = 4.12, p < 0.001, h = 0.24) than expected by chance. **(C)** Cosine similarity was computed between all pairs of synchronous calcium events (SCEs) within each session based on the overlap of active neurons. Distributions were generated either from all SCEs (“ALL”) or from the subset of SCEs involving at least one tdTomato-positive iEngram neuron (“RED”). During the A-Late phase, RED SCEs showed significantly higher population similarity than the full set of SCEs (Mann-Whitney U = 1678, p < 0.001, rank-biserial correlation = −0.713), indicating more structured and stereotyped ensemble recruitment. No difference was observed in non-learners (Mann-Whitney U = 3468, p = 0.396, rank-biserial correlation = −0.927).

To interpret how neural dynamics relate to successful behavioral recovery, we first examined whether memory reinstatement was consistent across animals or reflected distinct behavioral subgroups. Behavioral analysis revealed clear variability within the P19-trained group, with statistical tests supporting a bimodal distribution of freezing scores (Fig. S13A). Animals segregated into two groups: one with freezing levels comparable to naϊve controls (“non-learners”) and another with robust freezing responses, indicating successful reinstatement of the infantile memory (“learners”). This classification provided a basis for comparing neural ensemble dynamics between learners and non-learners to determine how SCE activity during A-Late relates to memory reinstatement.

Learners displayed a higher frequency of SCEs at any time during offline recordings (Fig. S13B), and their SCEs engaged a larger fraction of the CA3 network than non-learners (Fig. S13C). Independent from the ensembles they belonged to, individual neuron participation in SCEs positively correlated with overall activity rates during offline periods (Fig. S13D-F). As expected, CONTEXT and AVERSIVE ensemble neurons were preferentially recruited into SCEs during the Early and Late offline periods following their respective reminders (Fig. S13D-F, left). In contrast, iEngram and REINSTATED neurons were selectively recruited into SCEs during the Late offline period after the AVERSIVE reminder (Fig. S13D-F, right), suggesting this time window might be critical for any interaction between the two populations. To explore these interactions, we quantified the probability that iEngram neurons co-activated with other ensembles during SCEs, and tested against shuffled controls. While overall co-activation probabilities were generally low and population effects were negligible (Fig. S14A and B), the REINSTATED ensemble uniquely showed significantly elevated co-activation with iEngram neurons during SCEs taking place specifically during A-Late (Fig. S14A and B), and selectively in learners (Fig. 5B and S14A and B). Cosine similarity analysis revealed that SCEs involving iEngram neurons during this period were more similar to one another than SCEs in the overall population (Fig. 5C and S14C), indicating that iEngram-associated events occur in a more stereotyped network state. Together, these results demonstrate that successful reinstatement of a forgotten infantile memory is associated with selective and repeated co-activation of iEngram neurons with newly emerging CA3 ensembles during offline network-wide synchrony events. This structured interaction may provide the network-level mechanism by which a latent memory trace becomes integrated into a new ensemble supporting the reinstated memory recall.

## Discussion

The question of how seemingly inaccessible memories associated with early-life events continue to shape cognitive processes throughout life has been a central challenge in neuroscience and clinical psychology for over a century. In rodents, aversive infantile memories persist in a latent form in the adult brain, and under appropriate experimental circumstances, they can be reinstated and shape further learning. Here, we reveal that the reinstatement of a forgotten infantile memory in rodents requires a temporally structured sequence of distinct reminders and involves a carefully orchestrated hippocampus-centered network process. We propose that this process unfolds in three stages. First, a contextual reminder of the original experience primes the hippocampal network, enhancing the recruitment of neurons associated with the latent infantile memory (iEngram) upon subsequent stimulation (“Priming”, Fig. S15A). Next, a reminder of the aversive infant experience tags iEngram neurons for offline reactivation in the following hours (“Tagging”, Fig. S15B). Finally, heightened iEngram activity during offline reactivation events binds the latent infantile memory with novel neuronal ensembles, likely through synchronous network-wide events, thereby reinstating behavioral expression of the original memory (“Binding”, Fig. S15C).

These findings revise the traditional view of infantile memory engrams as silent placeholders of latent information. Instead, we show that hippocampal CA3 iEngram neurons remain embedded in adult network dynamics and actively contribute to memory processing across developmental stages. In fact, their transient recruitment to memory ensembles correlated with successful reinstatement processes, and their reactivation, during both the presentation of the aversive reminder and during subsequent offline periods, was necessary for the reinstatement of the latent memory. This evidence positions iEngram neurons as potential targets for interventions aimed at preventing unwanted reinstatement of aversive infantile memories.

Notably, the reinstated memory exhibited features typically associated with memories encoded during adulthood, such as specificity and persistence. Yet, the behavioral response it elicited remained consistent with the original early-life experience, suggesting that reinstatement involves more than the simple retrieval of a dormant memory trace. Instead, it appears to reflect a process whereby latent infantile representations are integrated into adult-like substrates. Supporting this view, the expression of the reinstated memory relied on the activation of a distinct neuronal ensemble emerging during a sensitive window following the presentation of the aversive reminder. Repeated co-activation between this novel ensemble and iEngram neurons in network-wide synchronous activity events at this time correlated with successful memory reinstatement. In contrast, disruption of iEngram activity, or the introduction of evidence of safety such as those underpinning extinction learning, prevented reinstatement. The existence of this critical window and the interplay between iEngram neurons and the surrounding network suggest that reinstatement involves a re-encoding process in which a latent memory becomes represented by a newly formed neuronal ensemble. Our data indicate that iEngram neurons act as an internal scaffold during offline network dynamics, guiding the formation of novel memory ensembles while anchoring them to early-life experiences. This mechanism represents a departure from classical models of memory retrieval or reconsolidation, pointing instead to a dynamic process in which latent traces are re-instantiated within novel neuronal substrates. These findings indicate that offline activity dynamics have the potential to shape the architecture of memory across the lifespan, and might open a window for potential non-invasive interventions to prevent the maladaptive reinstatement of latent memories arising from early-life traumatic events.

---

**Fig. S1.**
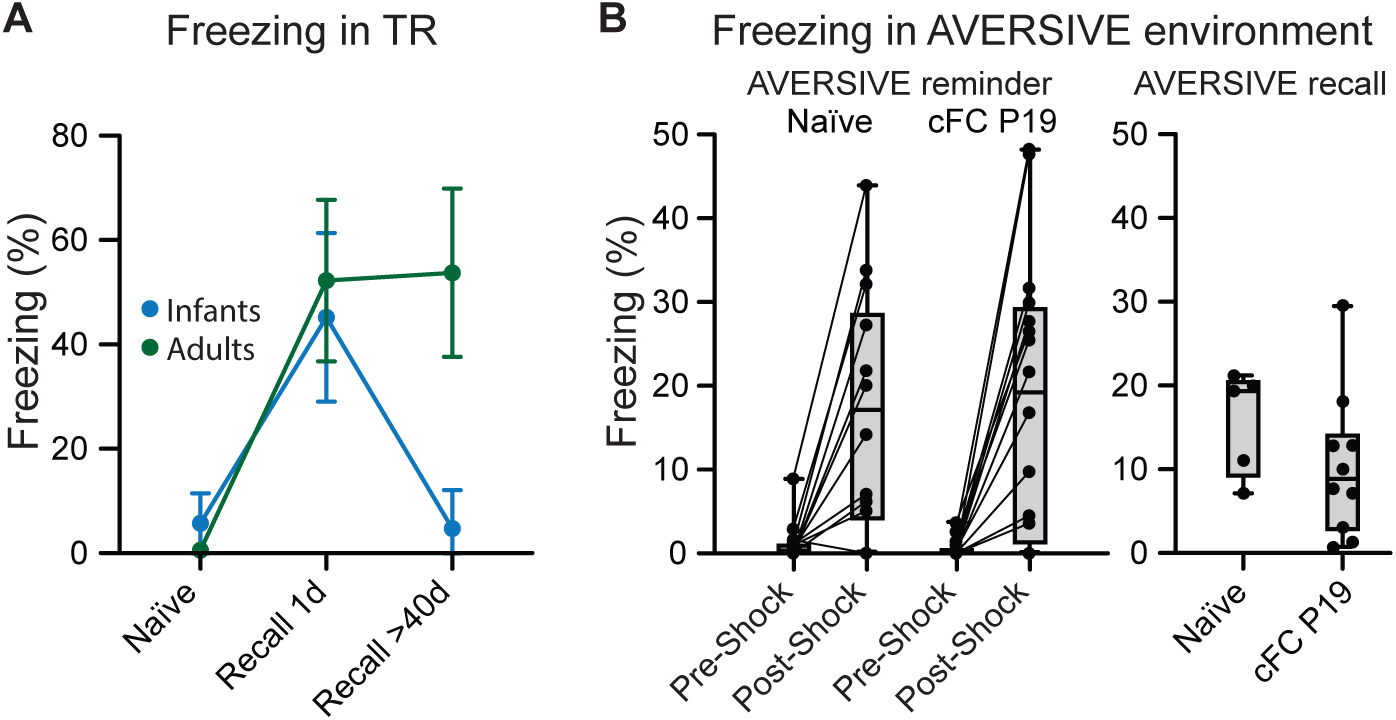
Infant cFC learning is forgotten at remote time points, leading to naϊve-like responses to AVERSIVE stimuli in adulthood. **(A)** Freezing responses in TR at recent (1 day) and remote (> 40 days) recall following cFC in infants (P19) or in adults (> P60). Mice conditioned at P19 showed robust freezing at recent recall, comparable to adult-trained animals. However, at remote recall, P19-conditioned mice froze at naϊve levels, indicating infantile amnesia, whereas adult-conditioned animals maintained elevated freezing responses. Repeated-measures two-way ANOVA with Geisser-Greenhouse correction: session x age interaction, F(1.658, 31.49) = 67.45, p < 0.001, 112 = 0.78; session, F(1.763, 44.07) = 194.6, p < 0.001, 112 = 0.886; age, F(1.000, 25.00) = 63.62, p < 0.001, 112 = 0.718. n ≥ 20 per group. **(B)** Freezing responses to an AVERSTVE shock in a novel context in adulthood did not differ between mice previously conditioned at P19 and naϊve animals (left). Two-way ANOVA: group, F(1, 28) = 0.01268, p = 0.9111; time, F(1, 28) = 38.36, p < 0.0001, 112 = 0.0006; interaction, F(1, 28) = 0.1236, p = 0.7277. Re-exposure to the environment where the AVERSTVE reminder occurred elicited low freezing levels in both naϊve and P19-conditioned animals (right). No significant difference between groups: t(13) = 1.230, p = 0.240, Cohen’s d = 0.104.

**Fig. S2.**
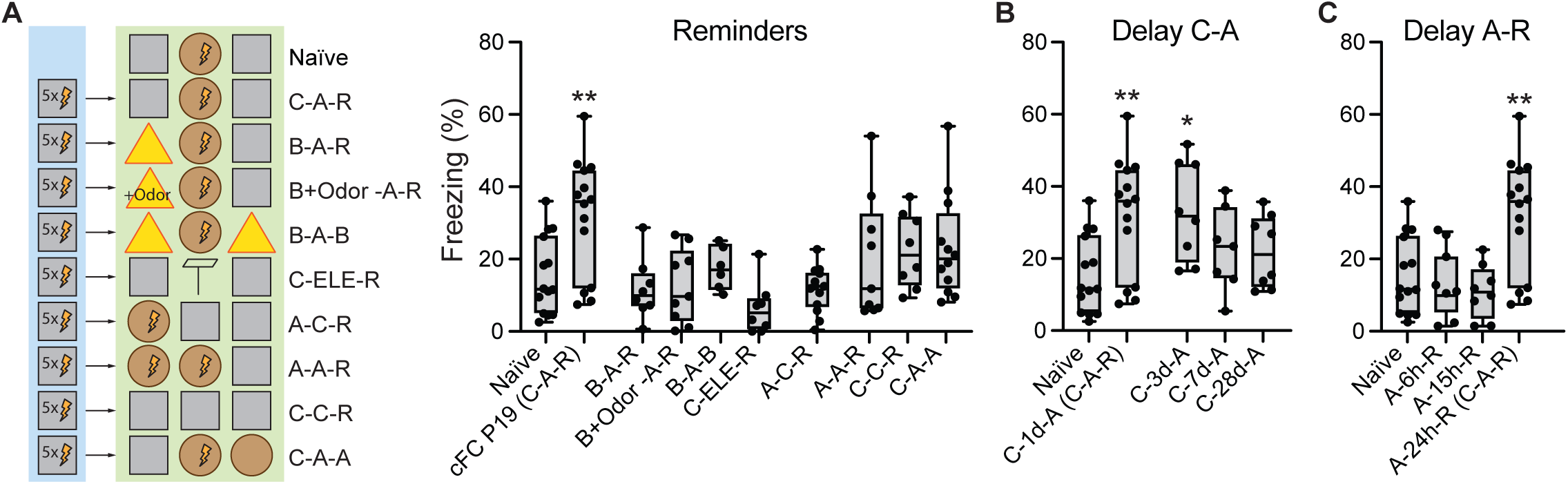
The reinstatement of a forgotten infantile memory relies on the timely and sequential presentation of CONTEXT and AVERSTVE reminders. **(A)** Right: Schematic representations of varying protocols for memory reinstatement. Left: Reinstatement required the specific presentation of a CONTEXT reminder (C) followed by a reminder of the specific AVERSIVE (A) stimulus experienced during cFC training at P19 (“Naϊve C-A-R” and “cFC P19 C-A-R”, as in Fig. 1C). Substituting the CONTEXT reminder (C) with a novel context B (“B-A-R” or “B-A-B”), pairing context B with a TR-matching odor (“B+Odor -A-R”), or replacing the AVERSIVE experience with exposure to an elevated platform (“C-ELE-R”) failed to induce freezing at REINSTATED recall (Dunnett’s test: B-A-R, p > 0.999; B-A-B, p > 0.999; B+Odor -A-R, p > 0.999; C-ELE-R, p = 0.756). Likewise, changing the order of reminders (“A-C-R”), or repeating a single reminder (“A-A-R” or “C-C-R”), did not reinstate memory (all p > 0.75). Only the canonical C-A-R condition produced high freezing. Returning to the AVERSIVE environment instead of TR at recall (“C-A-A”) did not elicit reinstatement (p = 0.807). One-way ANOVA: F(25, 226) = 11.18, p < 0.001, 12 = 0.5529; Brown-Forsythe: F(25, 226) = 2.622, p < 0.001; Bartlett’s test: x2 = 81.17, p < 0.001; n 2 6 per group. **(B)** Delays between CONTEXT and AVERSIVE reminders up to 3 days still triggered reinstatement (C-3d-A: p = 0.024), whereas longer intervals failed to promote it (C-7d-A: p = 0.635, C-28d-A: p = 0.187). Freezing levels at recall in all effective conditions were significantly different from naϊve controls (p = 0.007-0.024). n 2 7 per group. **(C)** Varying the delay between AVERSIVE and REINSTATED recall revealed that reinstated memory expression requires a 24 h consolidation interval. At shorter delays (6 h and 15 h), mice failed to express freezing upon presentation of the TR (REINSTATED 6 h and 15 h: p > 0.999). n 2 8 per group.

**Fig. S3.**
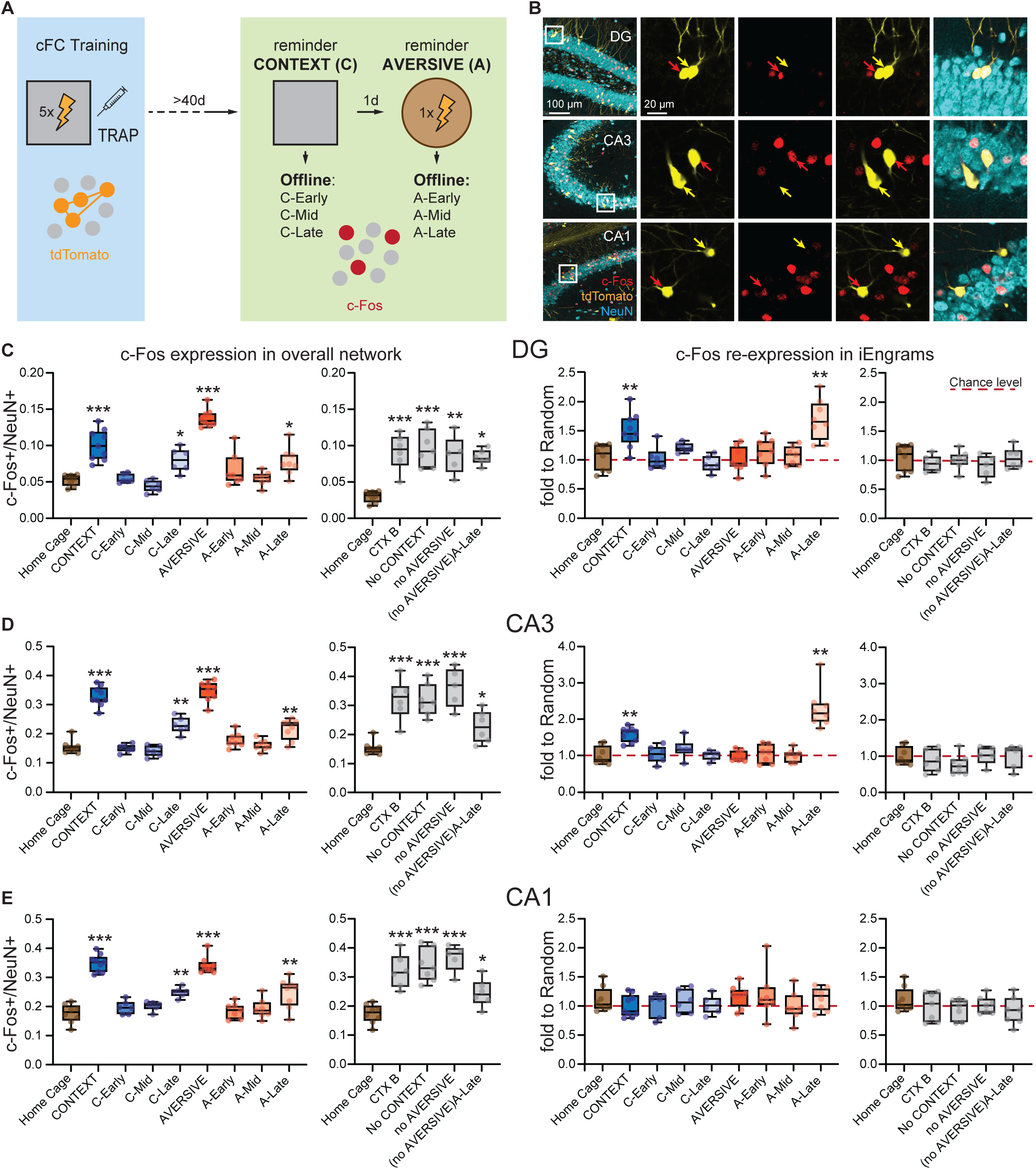
Recruitment of iEngram neurons in c-Fos+ ensembles throughout reinstatement. **(A)** Schematic of the experimental protocol. iEngram neurons were labeled in infancy via TRAP (Fos-driven tdTomato expression) during cFC training at P19. In adulthood, animals were perfused after specific behavioral sessions of the reinstatement protocol, including CONTEXT and AVERSIVE reminders, or during offline phases: Early (+3-5 h), Mid (+7-9 h), and Late (+13-15 h) after each reminder. **(B)** Representative images showing tdTomato (yellow), c-Fos (red), and NeuN (cyan) immunolabeling in DG (Top row), CA3 (Middle row), and CA1 (Bottom row) (maximum intensity projection, 30x objective, scale bar = 100 µm). Insets show single-plane images with yellow arrows indicating tdTomato+ c-Fos-neurons, and red arrows indicating tdTomato+ c-Fos+ neurons (scale bar = 20 µm). **(C)** In the dentate gyrus (DG), overall network c-Fos expression (c-Fos+/NeuN+, left) was elevated after CONTEXT and AVERSIVE reminders and during respective Late offline periods, as well as in control sessions associated to protocols that failed to trigger reinstatement (one-way ANOVA, F(12, 76) = 14.34, p < 0.001, 12 = 0.693). However, c-Fos re-expression in iEngram neurons (expressed as fold over random, right) increased only at CONTEXT and A-Late timepoints. **(D)** Similar trends were observed in CA3: increased c-Fos levels after CONTEXT, AVERSIVE, C-Late, and A-Late phases, and during non-reinstating control conditions (one-way ANOVA, F(12, 76) = 29.44, p < 0.001, 12 = 0.823). c-Fos re-expression in iEngram neurons was elevated at CONTEXT and A-Late only, mirroring DG. **(E)** In CA1, while overall c-Fos expression was modulated similarly to DG and CA3 (one-way ANOVA, F(12, 76) = 23.71, p < 0.001, 12 = 0.789), no increase in c-Fos re-expression was observed in iEngram neurons at any timepoint, suggesting subregion-specific recruitment dynamics.

**Fig. S4.**
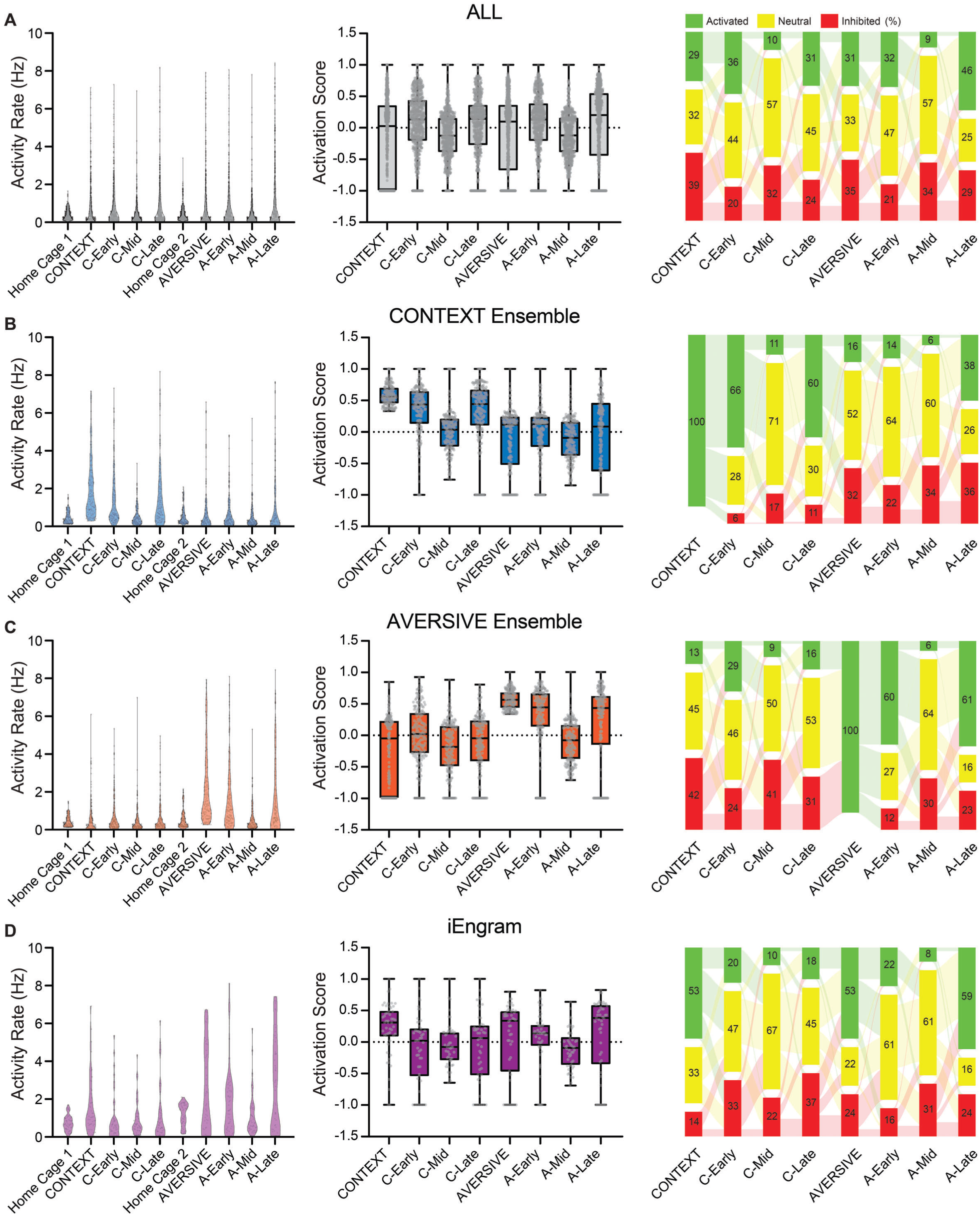
Activity dynamics of longitudinally tracked neurons across reminder sessions and subsequent offline phases. **(A-D)** Activity rates and tuning scores of longitudinally tracked neurons in the CA3 pyramidal layer across all behavioral sessions and offline phases. **(A)** All neurons (n = 690; 7 mice). Left: Activity rates. Friedman test revealed significant session-dependent modulation (x2(9) = 221.2, p < 0.001, W = 0.221). Post hoc Dunn’s comparisons showed increased activity at CONTEXT (p < 0.001), C-Early (p < 0.001), C-Late (p < 0.001), AVERSIVE (p < 0.001), A-Early (p < 0.001), and A-Late (p < 0.001), while C-Mid (p = 0.87), A-Mid (p = 0.111), and Home Cage 2 (p = 0.102) were not significantly different from baseline (Home Cage 1). Center: AS varied significantly across sessions (repeated-measures one-way ANOVA with Geisser-Greenhouse correction: F(5.503, 3792) = 25.34, p < 0.001, f2_partial = 0.036, s = 0.7862). Two-tailed one-sample Wilcoxon Signed Rank Test vs. baseline activation: CONTEXT (W = −61066, p < 0.0001), C-Early (W = 57389, p < 0.0001), C-Mid (W = −803, p = 0.128), C-Late (W = 32972, p = 0.0016), AVERSIVE (W = −35114, p = 0.0008), A-Early (W = 42688, p < 0.0001), A-Mid (W = −739, p = 0.078), A-Late (W = 18325, p = 0.0396). Right: neurons were categorized as activated (AS > 0.33), neutral −0.33 :S AS :S 0.33), or inhibited (AS < −0.33). Sankey plots depict the transitions between activity states across sessions, revealing dynamic recruitment patterns during both reminders and the associated offline phases. **(B)** CONTEXT ensemble (n = 184; 7 mice). Left: Activity rates significantly varied across sessions (Friedman test, x2(9) = 448.6, p < 0.001, W = 0.222). Increased activity was detected at CONTEXT (p < 0.0001), C-Early (p < 0.0001), and C-Late (p < 0.0001). No changes were observed at C-Mid (p > 0.9999), AVERSIVE (p > 0.9999), A-Early (p > 0.9999), A-Mid (p > 0.9999), A-Late (p = 0.5328), or Home Cage 2 (p > 0.9999). Center: AS dynamics reflected increased activation at CONTEXT, C-Early, and C-Late (Two-tailed one-sample Wilcoxon Signed Rank Test vs. baseline activation: CONTEXT, W = 17020; C-Early, W = 12897; C-Late, W = 10503; all p < 0.0001). No significant change at C-Mid (W = −224, p = 0.878), AVERSIVE (W = −2446, p = 0.091), A-Early (W = 855, p = 0.556), A-Mid (W = −6108, p = 0.084) or A-Late (W = −1931, p = 0.179). Repeated-measures one-way ANOVA with Geisser-Greenhouse correction: F(5.105, 934.2) = 69.81, p < 0.001, f2_partial = 0.276, s = 0.7293. Data reported as in Fig. 2E. Right: neurons were categorized as activated (AS > 0.33), neutral −0.33 :S AS :S 0.33), or inhibited (AS < −0.33). Sankey plots depict the transitions between activity states across sessions, revealing dynamic recruitment patterns during both reminders and the associated offline phases. **(C)** AVERSIVE ensemble (n = 195; 7 mice). Left: Activity rates varied significantly across sessions (Friedman test, x2(9) = 506.7, p < 0.001, W = 0.236). Elevated activity was observed at AVERSIVE (p < 0.0001), A-Early (p < 0.0001), and A-Late (p < 0.0001). No significant differences from baseline were found at CONTEXT (p = 0.2661), C-Early (p > 0.9999), C-Mid (p = 0.35), C-Late (p > 0.9999), or Home Cage 2 (p > 0.9999). Center: AS dynamics showed increased activity during AVERSIVE (W = 19110, p < 0.001), A-Early (W = 10956, p < 0.0001), and A-Late (W = 4009, p < 0.001), but not during CONTEXT (W = −7362, p = 0.327), C-Early (W = 564, p = 0.722), C-Mid (W = −8976, p = 0.083), C-Late (W = −3870, p = 0.074), or A-Mid (W = −6328, p = 0.092). Repeated-measures one-way ANOVA with Geisser-Greenhouse correction: F(5.011, 972.1) = 71.80, p < 0.001, f2_partial = 0.270, s = 0.7158. Data reported as in Fig. 2E. Right: neurons were categorized as activated (AS > 0.33), neutral −0.33 :S AS :S 0.33), or inhibited (AS < −0.33). Sankey plots depict the transitions between activity states across sessions, revealing dynamic recruitment patterns during both reminders and the associated offline phases. **(D)** iEngram neurons (n = 50; 7 mice). Left: Activity dynamics were also session-specific (Friedman test, x2(9) = 221.2, p < 0.001, W = 0.151). Significant increases occurred at CONTEXT (p = 0.0075), AVERSIVE (p = 0.0064), and A-Late (p = 0.0001). Other sessions were not different from baseline: C-Early (p > 0.9999), C-Mid (p > 0.9999), C-Late (p > 0.9999), A-Early (p = 0.077), A-Mid (p = 0.9406), and Home Cage 2 (p = 0.229). Center: AS dynamics reflected significantly activated neurons during CONTEXT (W = 690, p < 0.001), AVERSIVE (W = 140, p = 0.0381), and A-Late (W = 229, p = 0.0141), but not during C-Early (W = −231, p = 0.2681), C-Mid (W = −239, p = 0.2531), A-Early (W = 429, p = 0.504), or A-Mid (W = −505, p = 0.2727). Repeated-measures one-way ANOVA with Geisser-Greenhouse correction: F(5.345, 261.9) = 2.909, p = 0.012, f2_partial = 0.215, s = 0.7636. Data reported as in Fig. 2E. Right: neurons were categorized as activated (AS > 0.33), neutral −0.33 :S AS :S 0.33), or inhibited (AS < −0.33). Sankey plots depict the transitions between activity states across sessions, revealing dynamic recruitment patterns during both reminders and the associated offline phases.

**Figure S5.**
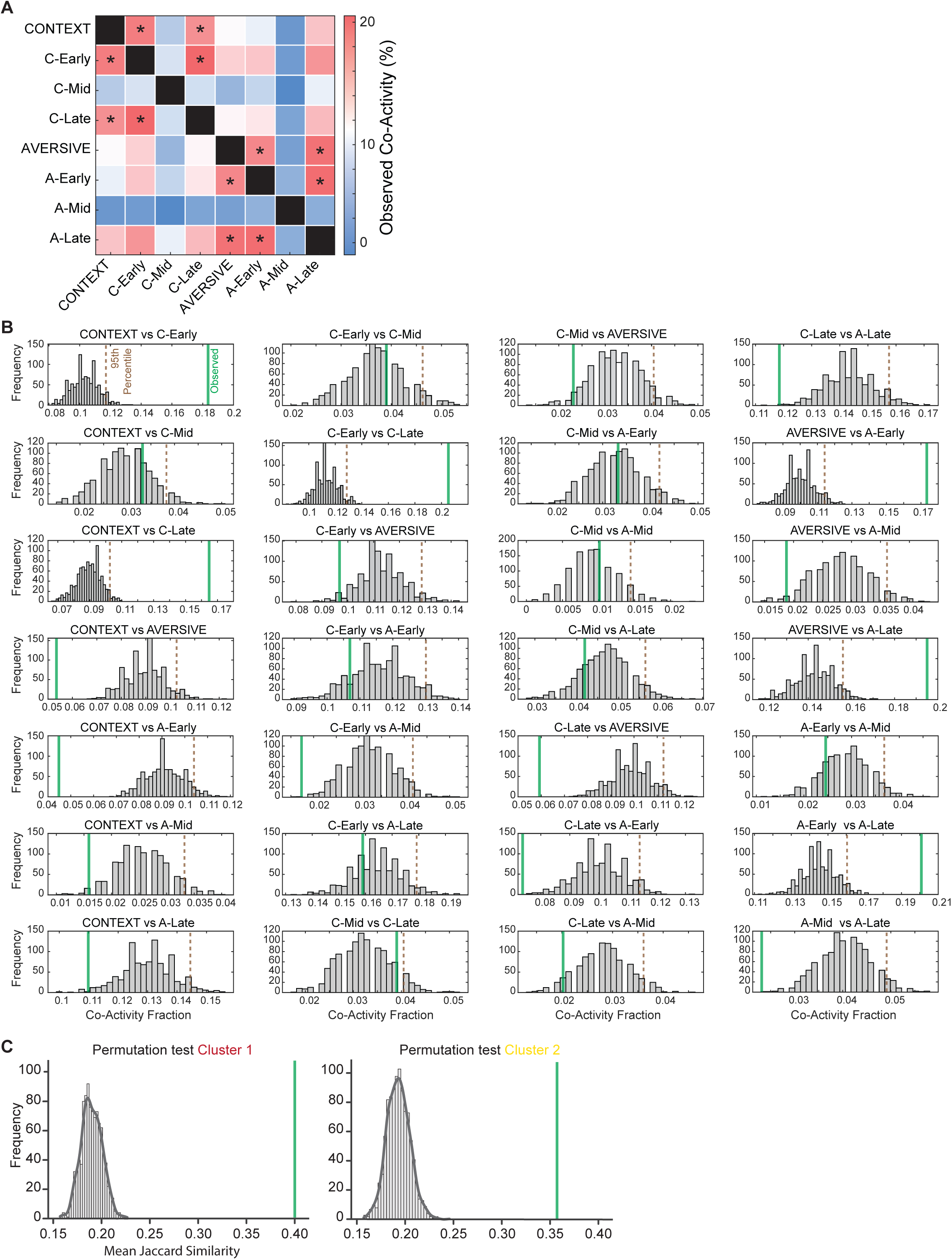
Permutation-based test of session pairwise co-activity overlap. **(A)** Matrix summarizing the significance of co-activity overlap between pairs of sessions. For each session pair, we quantified the fraction of neurons co-active in both conditions and assessed significance using a permutation-based method (1000 shuffles), comparing the observed overlap to the 95th percentile of the null distribution. Asterisks indicate comparisons in which the observed overlap exceeded the significance threshold (*p < 0.05). **(B)** Histograms showing the null distributions of co-activity overlap for all session pairs tested in (A). Green vertical lines mark the observed co-activity values used to determine statistical significance compared to 95th percentile (stapled brown line). **(C)** Permutation histograms of Jaccard similarity for condition-pair comparisons based on ensemble clustering. As in (B), green vertical lines represent observed values, and null distributions reflect shuffled condition identities.

**Fig. S6.**
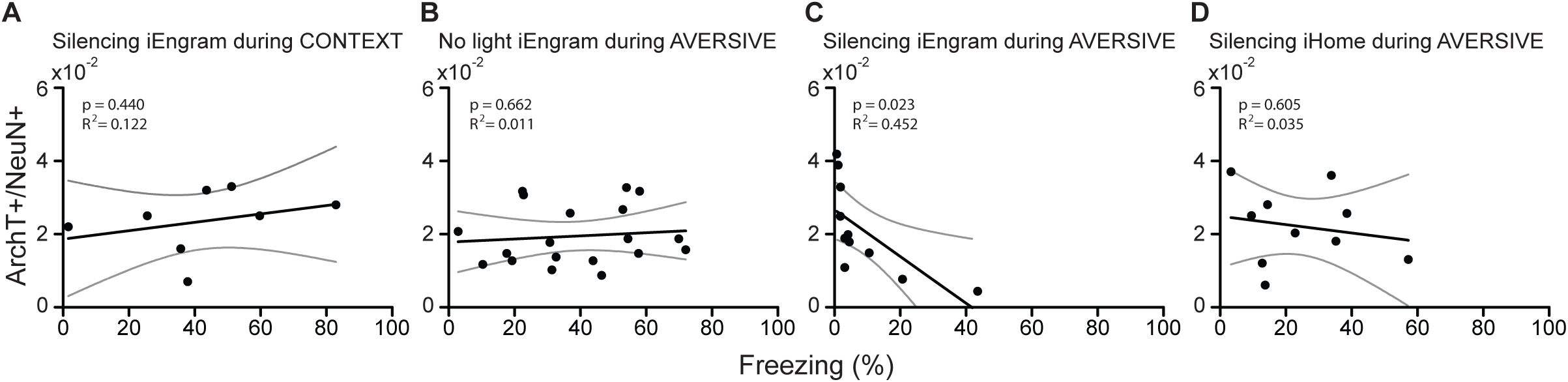
REINSTATED memory recall strength correlates with the fraction of targeted iEngram neurons during optogenetic silencing at the AVERSIVE reminder. **(A)** A simple linear regression was used to assess whether the fraction of CA3 pyramidal neurons expressing ArchT (ArchT+/NeuN+) predicted freezing behavior during REINSTATED recall upon iEngram optogenetic silencing. No significant correlation was observed when iEngram neurons were silenced during the CONTEXT reminder (n = 8 mice; F(1,6) = 0.683, p = 0.440, R2 = 0.122). **(B)** No correlation was found in control animals that expressed ArchT in iEngram neurons but did not receive light stimulation during the AVERSIVE reminder (n = 19 mice; F(1,17) = 0.198, p = 0.662, R2 = 0.011). **(C)** When iEngram neurons were silenced during the AVERSIVE reminder, the fraction of ArchT-expressing neurons negatively predicted freezing behavior at REINSTATED recall (n = 11 mice; F(1,9) = 7.439, p = 0.023, R2 = 0.452). **(D)** Silencing neurons TRAPed in the home cage at P21 following cFC training at P19 (iHome) during the AVERSIVE reminder did not yield a significant relationship between the fraction of CA3 pyramidal neurons expressing ArchT (ArchT+/NeuN+) and freezing behavior during REINSTATED recall (n = 10 mice; F(1,8) = 0.291, p = 0.605, R2 = 0.035).

**Fig. S7.**
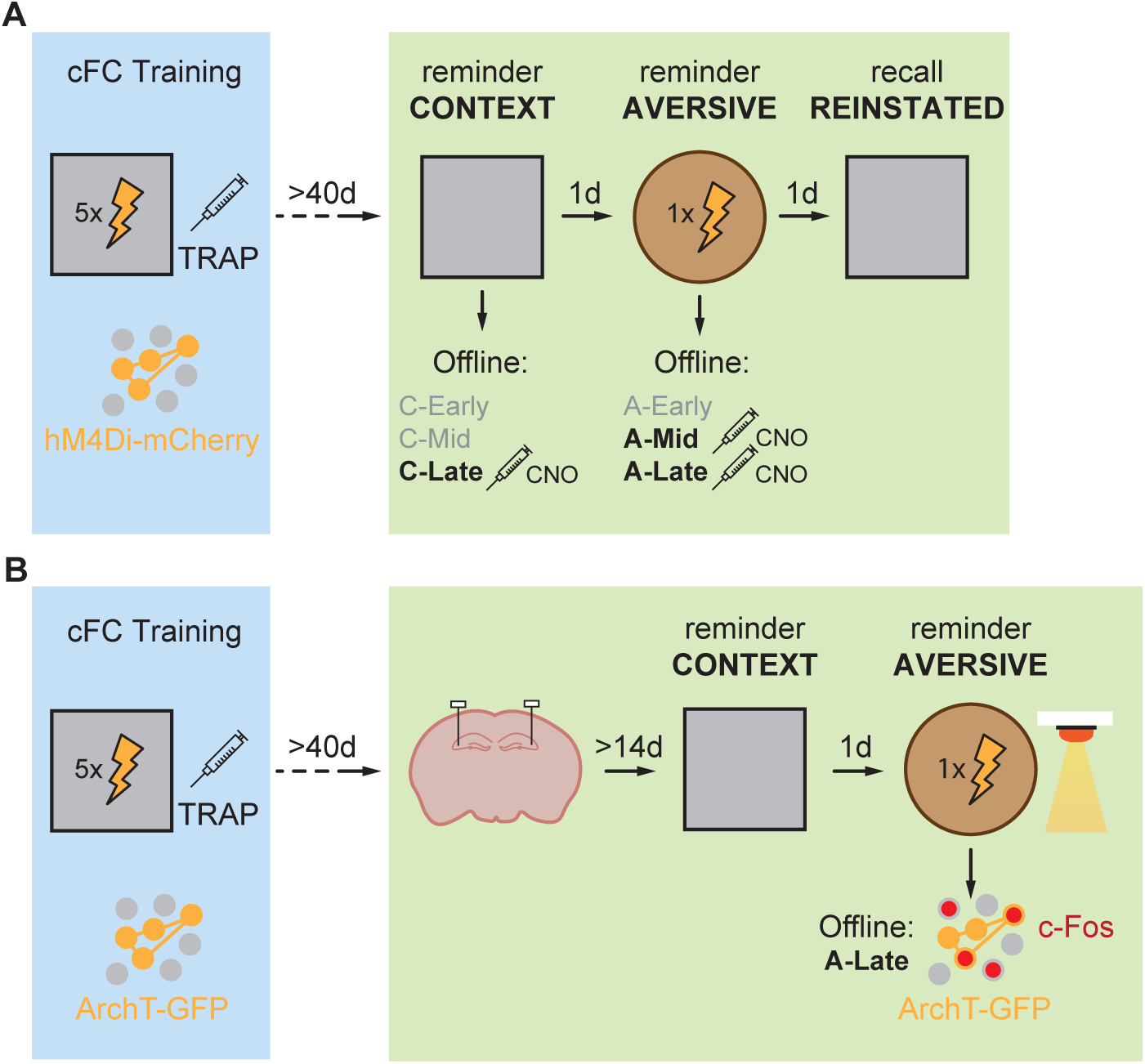
Memory reinstatement requires active iEngrams during A-late offline phase. **(A)** Schematic of the experiment for chemogenetic inhibition of iEngram neurons during offline phases of memory reinstatement. Expression of hM4Di-mCherry was induced in iEngram neurons during infant cFC at Pl9 using the TRAP system, followed in adulthood by CNO-mediated inhibition during discrete offline windows: C-Late, A-Mid, and A-Late. **(B)** Schematic of the protocol for optogenetic inhibition of iEngrams during the AVERSTVE reminder, followed by perfusion and c-Fos immunohistochemistry at the A-Late offline phase.

**Fig. S8.**
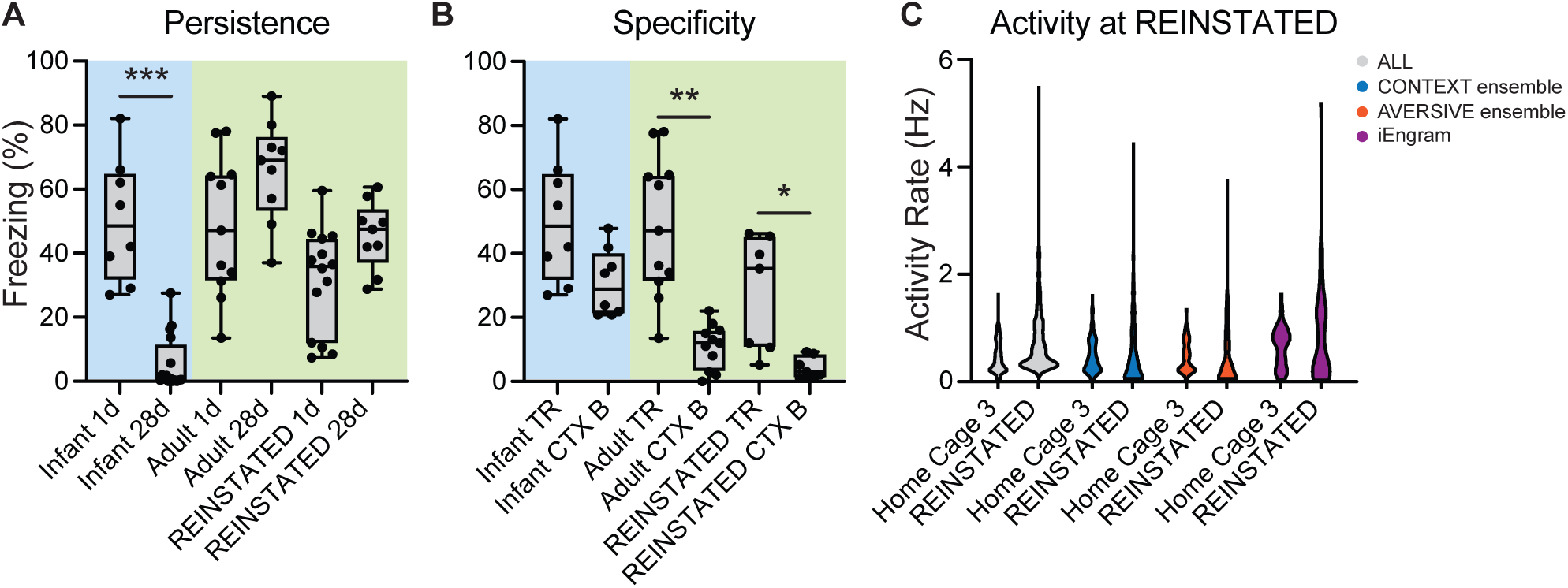
Reinstated memories display adult-like properties and do not rely on iEngram activity for their expression. **(A)** Infant cFC memories were expressed at recent recall (1d) but not at remote recall (28d), indicating infantile amnesia (n ≥ 8; two-tailed Mann-Whitney test: U = 1, p < 0.001, r = −0.788). In contrast, adult cFC memories were retained over time (n ≥ 9; U = 24, p = 0.562, r = −0.434), with a non-statistically significant trend towards higher freezing at longer delays. Similarly, reinstated memories showed robust expression at both recent and remote timepoints (n ≥ 9; U = 29, p = 0.330, r = 0.446), with a non-statistically significant trend towards higher freezing at longer delays. **(B)** One day following memory encoding, infant learners showed generalized fear during recall, with no significant difference in freezing between the training context (TR) and a novel context B (n = 8; Wilcoxon matched-pairs signed-rank test: W = −26.0, p = 0.078, r = 0.626). In contrast, at similar delays from encoding, adult learners exhibited context-specific memory recall (n = 11; W = −66.0, p = 0.001, r = 0.992). Reinstated memories also displayed context specificity, with significantly more freezing in the training context TR than in a novel context B (n = 7; W = −28.0, p = 0.016, r = 0.918). **(C)** Activity rates of neurons in hippocampal CA3 were compared between the REINSTATED recall session and the preceding home cage session (Home Cage 3). Significant differences were found when considering all tracked neurons together (n = 690 from 7 animals, two-tailed Wilcoxon matched-pairs signed rank test; W = 7465, p = 0.0087, r = 0.169), but no difference was found for CONTEXT (n = 184; W = −1888, p = 0.193, r = 0.096), AVERSIVE (n = 195; W = −2040, p = 0.197, r = 0.092), or iEngram neurons (n = 50; W = 117, p = 0.579, r = 0.078).

**Fig. S9.**
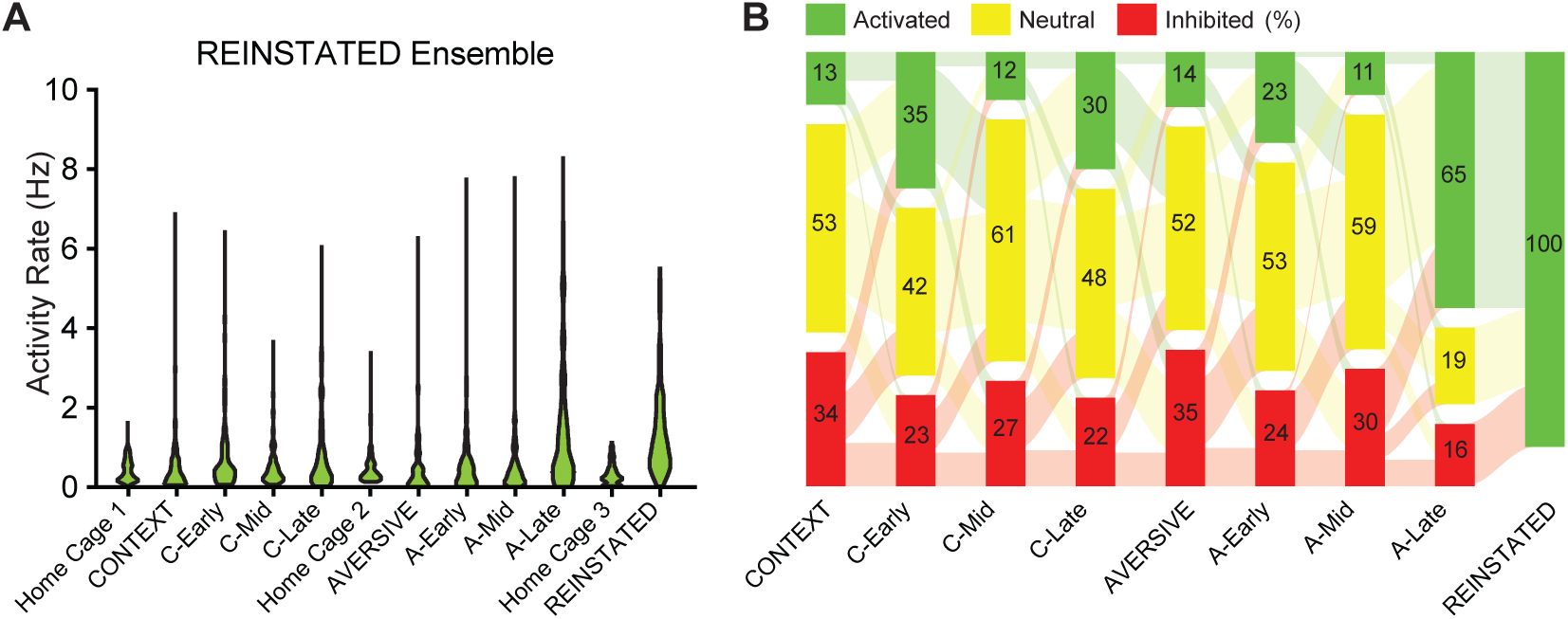
REINSTATED ensemble is highly active during A-late offline phase. **(A)** Activity rates of REINSTATED ensemble neurons (166 neurons from 7 mice) were compared across all sessions of memory reinstatement using a Friedman test (x2(11) = 290.8, p < 0.001, W = 0.1593). Dunn’s post hoc comparisons against Home Cage 1 revealed significantly elevated activity during A-late (p < 0.0001) and REINSTATED recall (p < 0.0001), but not in any other session (all p > 0.06). **(B)** REINSTATED ensemble neurons were categorized as activated (AS > 0.33), neutral −0.33 :S AS :S 0.33), or inhibited (AS < −0.33) based on their AS. Sankey plots depict the transitions between activity states across sessions, revealing dynamic recruitment patterns during both reminders and the associated offline phases. A majority of REINSTATED ensemble neurons were reactivated from A-late to REINSTATED recall, suggesting persistent engagement across this offline-to-recall transition.

**Figure S10.**
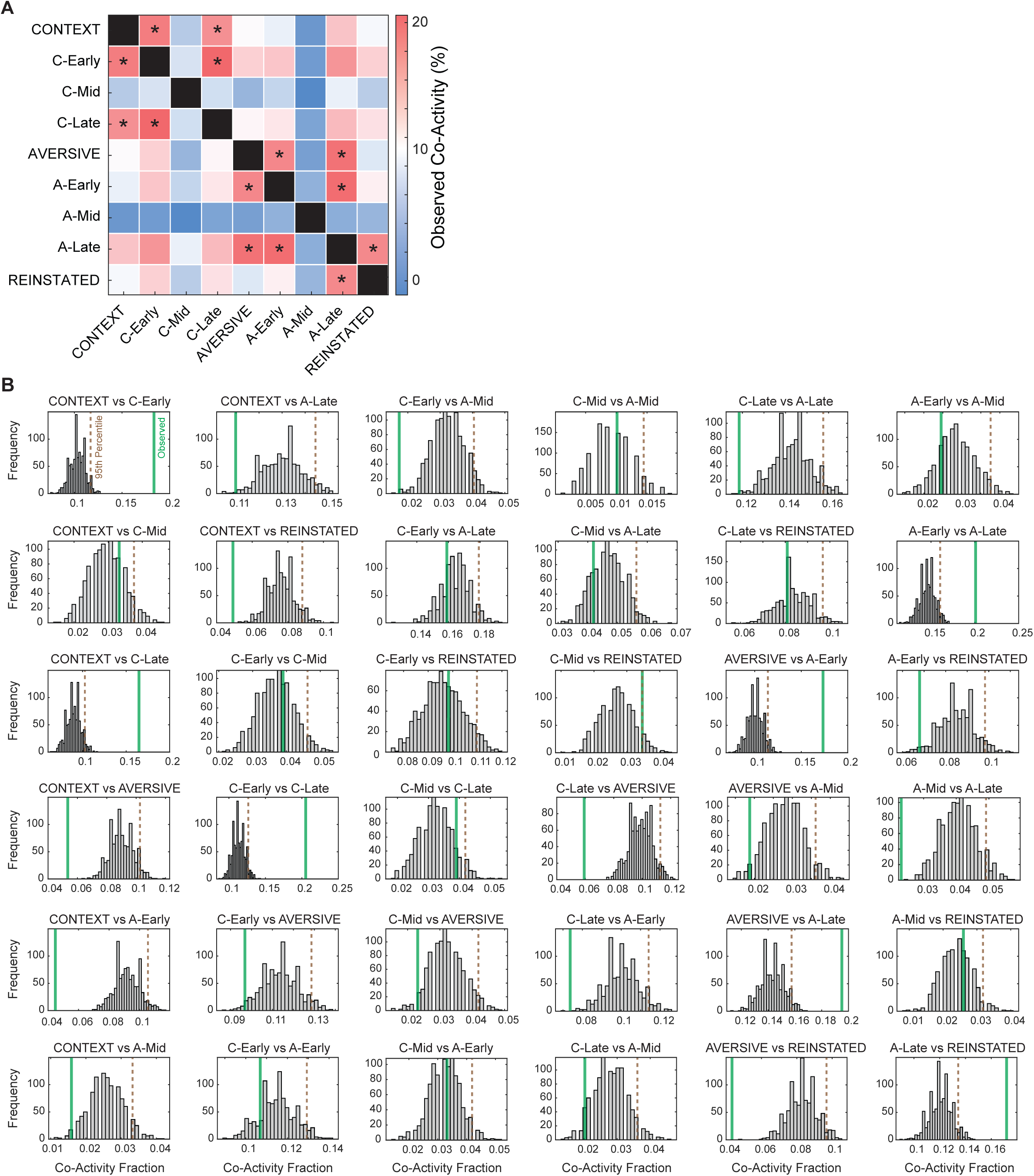
Permutation-based test of co-activity overlap with the REINSTATED ensemble. **(A)** Matrix summarizing the significance of neuronal co-activity overlap between pairs of sessions, now including the REINSTATED recall as an additional condition. For each pair, the fraction of neurons co-active in both sessions was compared against a null distribution generated by 1000 random permutations of session labels. Asterisks denote comparisons where the observed co-activity fraction exceeded the 95th percentile of the shuffled distribution (*p < 0.05). **(B)** Histograms of permutation-based null distributions for all pairwise comparisons shown in (A), including those involving the REINSTATED condition. Vertical lines indicate the observed co-activity values used to determine significance.

**Fig. S11.**
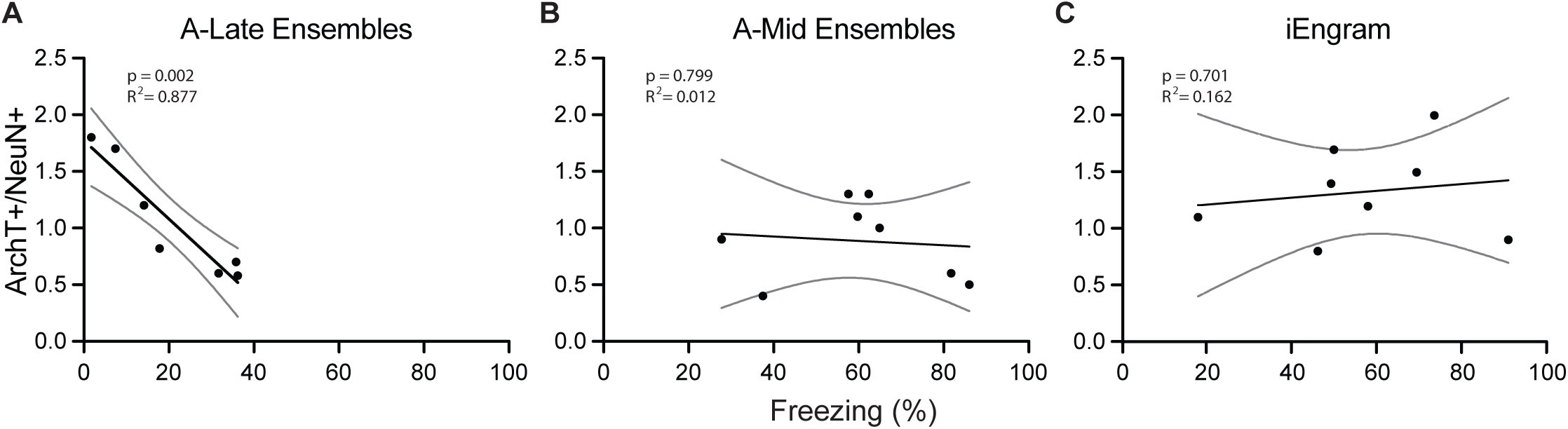
REINSTATED memory strength correlates with the fraction of TRAPed A-Late neurons during optogenetic silencing at recall. **(A)** A significant negative correlation was detected between freezing levels at RElNSTATED recall and the fraction of ArchT-expressing neurons (ArchT+/NeuN+) in hippocampal CA3 following optogenetic inhibition when A-Late ensemble neurons were TRAPed for ArchT expression (n = 7 mice; Simple linear regression; F(1,5) = 35.51, p = 0.002, R2 = 0.8772). **(B)** No correlation was found when inhibiting A-Mid ensemble neurons (n = 8; F(1,6) = 0.071, p = 0.799, R2 = 0.0122). **(C)** Similarly, no significant relationship was observed when iEngram neurons were targeted by expression of ArchT (n = 8; F(1,6) = 0.162, p = 0.701, R2 = 0.162).

**Figure S12.**
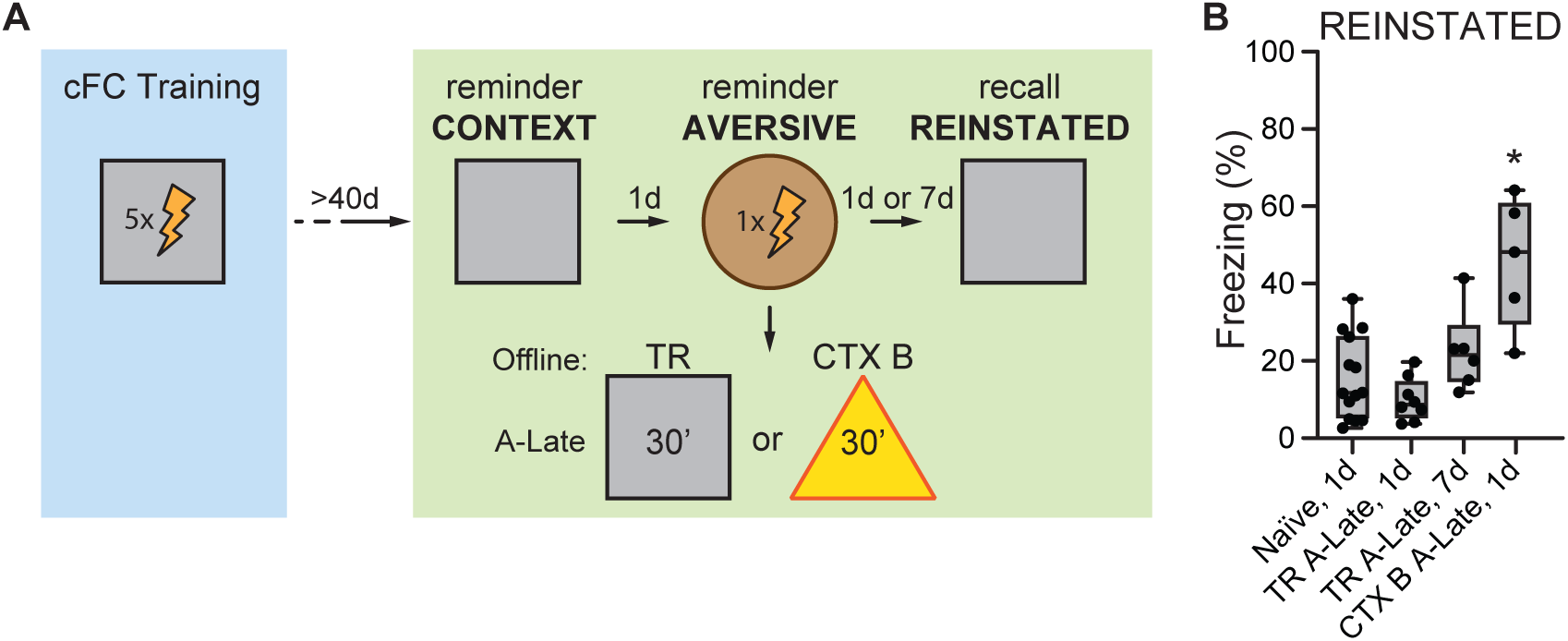
Reinstatement of infant fear memory is prevented by prolonged exposure to the training context during the A-Late offline period. **(A)** Schematic of the behavioral experiment. Mice were exposed for 30 minutes either to the training context (TR) or to a novel context (CTX B) during the A-Late period. Memory reinstatement was tested either 1 or 7 days later. **(B)** Mice exposed to the training context (TR) during the A-Late period froze to levels that were similar to those of naϊve animals when put back in TR during REINSTATED recall (n = 8, Kruskal Wallis test over all groups: H(4) = 14.59; p = 0.002, £2 = 0.4. Dunn’s multiple comparisons test to Naϊve for TR A-Late, 1d: p = 0.719). No difference compared to non-reinstating, naϊve animals was observed in animals that were tested for reinstatement at a distance of 7 days (n = 6, Dunn’s multiple comparisons test to Naϊve for TR A-Late, 7d: p = 0.434). Significantly high levels of freezing indicative of memory reinstatement were instead observed when animals were exposed to a novel context (CTX B) instead of TR during the behavioral intervention on the A-Late offline period (n = 5, Dunn’s multiple comparisons test to Naϊve for CTX B A-Late, 1d: p = 0.013).

**Figure S13.**
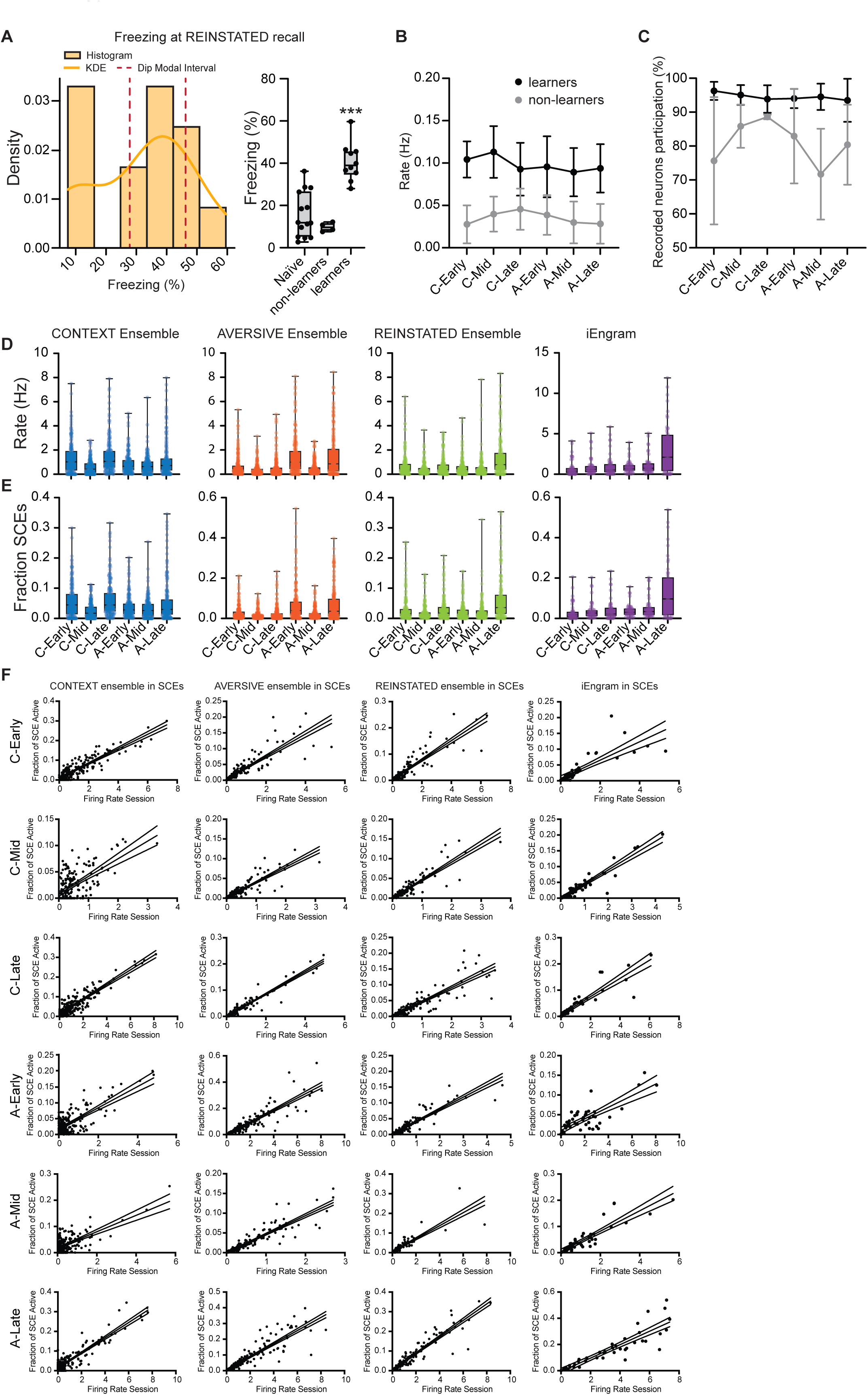
Increased synchronous calcium events in reinstating animals. **(A)** Left: histogram of freezing levels at REINSTATED recall. Gaussian mixture modeling (GMM) and modality testing (Hartigan’s dip test: D = 0.1105, p < 0.05; GMM ΔBIC = 10.3; 2 components favored in 90.1% of bootstraps) revealed a bimodal distribution. K-means clustering (k = 2) identified low (n = 4) and high (n = 10) freezing clusters. Right: high-freezing animals (“learners”) froze significantly more than both “non-learners” and naYve animals (Welch’s t-test: p < 0.001; Mann-Whitney U: p < 0.001); no significant difference between non-learners and naYve (Welch’s t-test: p = 0.0726; Mann-Whitney U: p = 0.5052). **(B)** Mice that reinstated the infant fear memory exhibited a significantly higher frequency of synchronous calcium events (SCEs) across all offline periods of the reinstatement protocol compared to non-learning animals. A two-way repeated measures ANOVA with Geisser-Greenhouse correction revealed a significant main effect of behavior (i.e. learners Vs. non-learners, F(1, 8) = 20.37, p = 0.002, 12 (partial) = 0.732). In contrast, the session and the interaction between session and behavior were not significant (F (2.697, 21.58) = 0.674, p = 0.563, 12 (partial) = 0.078; and F(2.697, 21.58) = 0.548, p = 0.637, 12 (partial) = 0.064, respectively). **(C)** Similarly, the average fraction of recorded neurons participating in all SCEs was significantly higher in reinstating animals compared to non-learners (main effect of behavior: F(1, 8) = 25.42, p = 0.001, 12_partial = 0.590), with no significant session or session x behavior interaction (F (2.139, 17.11) = 1.856, p = 0.185, 12 (partial) = 0.188; and F(2.139, 17.11) = 2.343, p = 0.123, 12 (partial) = 0.227, respectively). **(D, E)** The participation of neurons from four functionally defined neuronal ensembles (CONTEXT, AVERSIVE, REINSTATED, and iEngram) to SCEs was tracked across offline sessions following the CONTEXT and AVERSIVE reminders. All ensembles showed significant variation in participation to SCEs across sessions, quantified either by activity rates across SCEs frames (D) or fraction of SCEs frames in which any neuron was active (E), as assessed by Friedman tests (CONTEXT: x2(5) = 88.03, p < 0.001, W = 0.096; AVERSIVE: x2(5) = 141.5, p < 0.001, W = 0.145; REINSTATED: x2(5) = 75.43, p < 0.001, W = 0.091; iEngram: x2(5) = 22.74, p < 0.001, W = 0.090). **(F)** For all ensembles and offline periods, linear regression analyses revealed a strong positive relationship between a neuron’s average firing rate and the fraction of SCEs it participated in. All regressions were highly significant (p < 0.001) with large effect sizes (R2 ranging from 0.430 to 0.888).

**Figure S14.**
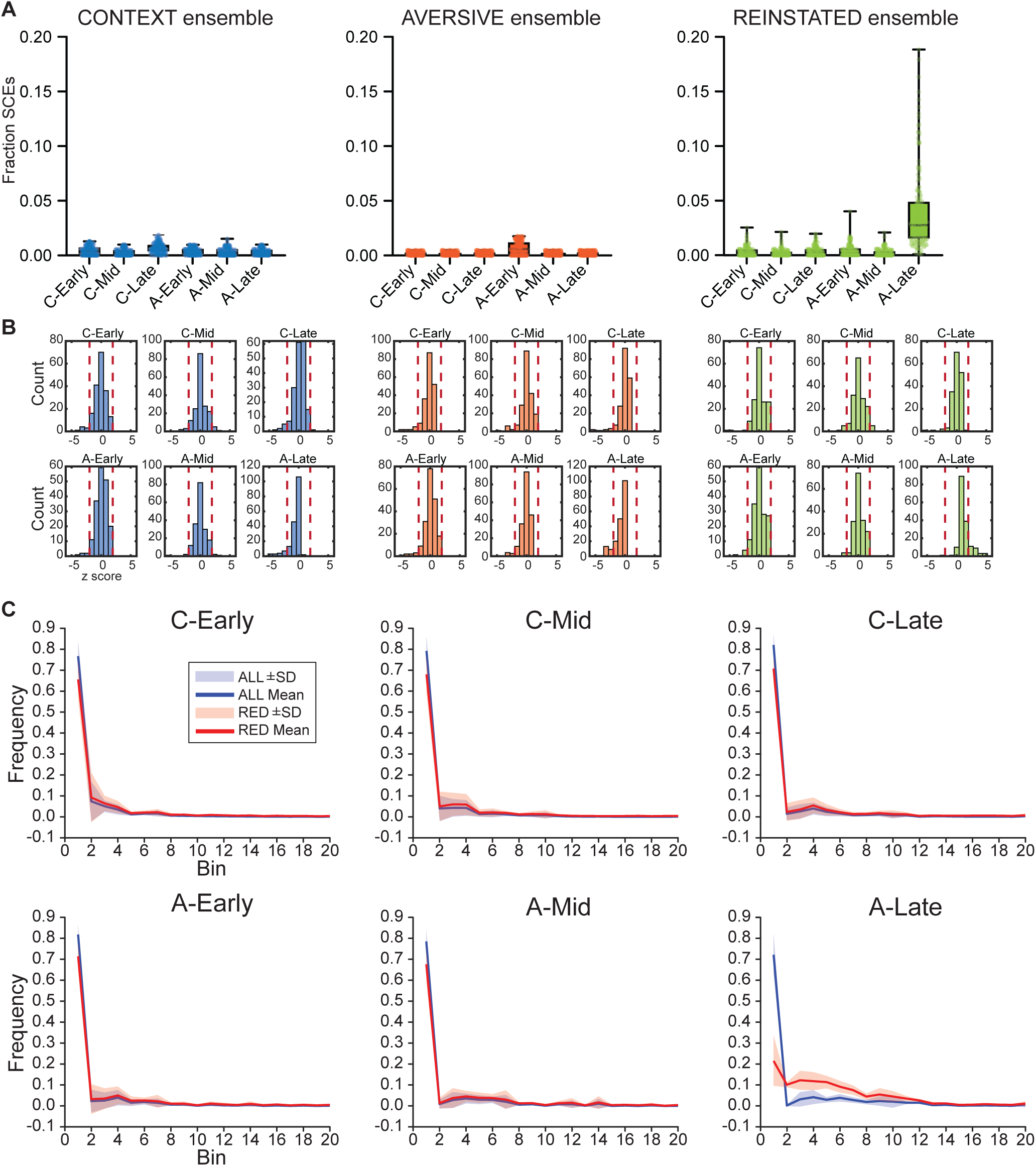
Selective ensemble coactivation and neuronal overrepresentation in iEngram-involving synchronous events for learners. **(A)** Fraction of synchronous calcium events (SCEs) that included at least one iEngram neuron and also recruited neurons from the CONTEXT, AVERSIVE, or REINSTATED ensembles, measured across CONTEXT (C-Early, C-Mid, C-Late) and AVERSIVE (A-Early, A-Mid, A-Late) offline sessions. All three ensembles showed significant variation in co-recruitment dynamics across sessions (Friedman tests: CONTEXT: x2(5) = 112.7, p < 0.001, W = 0.122; AVERSIVE: x2(5) = 54.88, p < 0.001, W = 0.056; REINSTATED: x2(5) = 345.4, p < 0.001, W = 0.041). While these shifts were significant, the associated effect sizes (Kendall’s W) were small, indicating that the magnitude of this modulation was mild. **(B)** Distribution of z-scores quantifying each individual neuron’s participation in SCEs, normalized against chance levels derived from firing rate-matched surrogates. Each subplot shows the session-specific distribution for neurons in a given ensemble. Red dashed lines indicate critical z-scores (±1.96), corresponding to the 0.05 significance level (two-tailed). Axes are fixed from −5 to +5 to enable visual comparison across conditions. **(C)** Similarity between synchronous calcium events (SCEs) was quantified using the cosine similarity measure. Each SCE was represented as a population vector, where individual vector elements corresponded to recorded neurons, and neuronal activity was encoded as binary values (1 = active, 0 = inactive). Cosine similarity was then calculated between all pairs of these vectors to quantify the overlap in active neuronal populations across SCEs. Frequency distributions of similarity measures were generated either from comparisons involving all SCEs (“ALL”); or from the subset of SCEs involving at least one tdTomato-positive iEngram neuron (“RED”). Across sessions, the similarity distributions largely overlapped; however, during the A-Late phase, RED SCEs showed significantly higher population similarity than the full set of SCEs (Mann-Whitney U = 1678, p < 0.001, rank-biserial correlation = −0.713), indicating more structured and stereotyped ensemble recruitment in iEngram+ SCEs (as reported in Fig. 5C). No significant differences were observed in earlier sessions (C-Early: U = 178, p = 0.097, r = −0.033; C-Mid: U = 259, p = 0.062, r = −0.328; C-Late: U = 401, p = 0.073, r = −0.210; A-Early: U = 249, p = 0.123, r = −0.015; A-Mid: U = 735, p = 0.061, r = −0.382).

**Figure S15.**
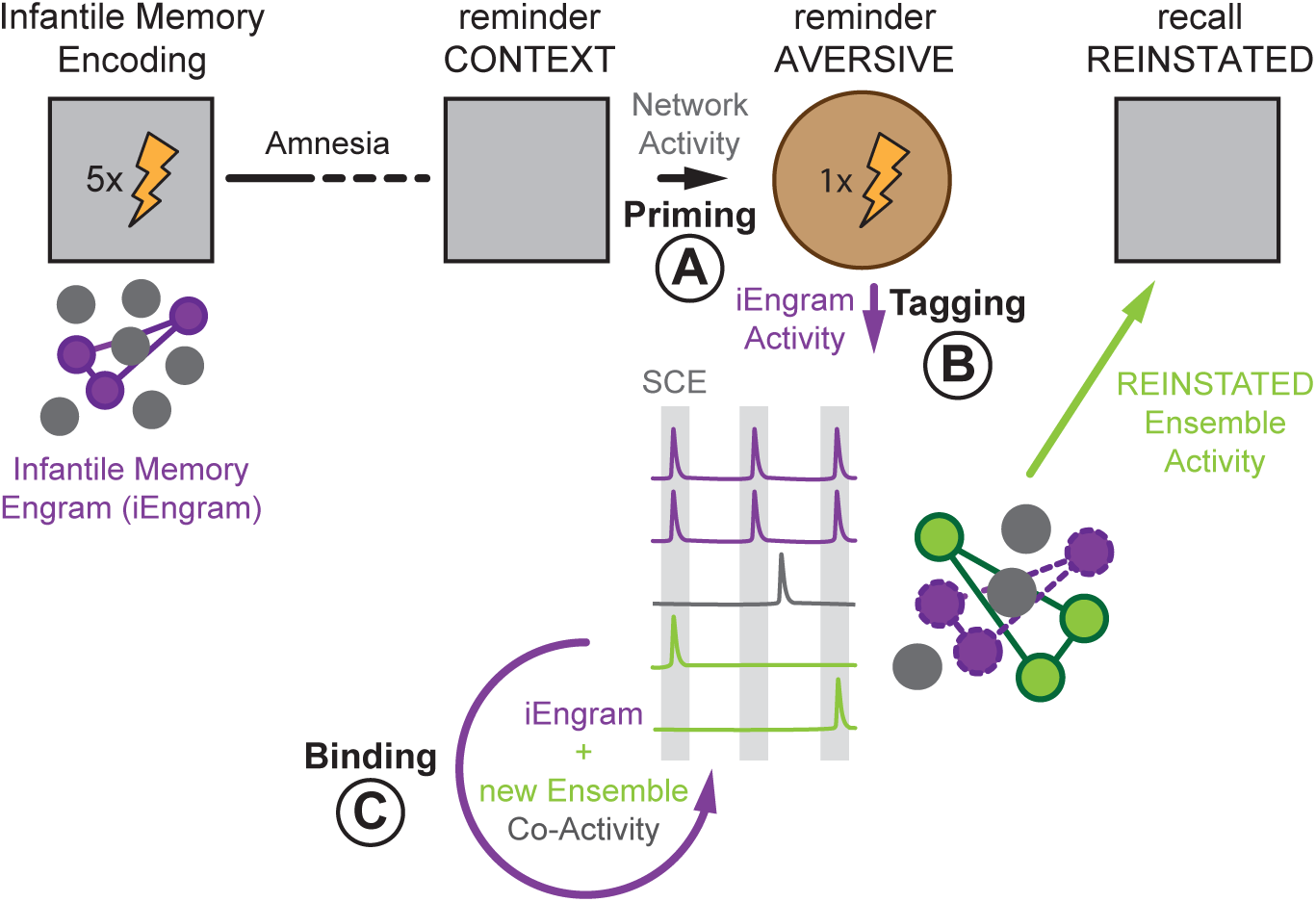
A network model for the reinstatement of a forgotten infantile memory. We propose that reinstating forgotten infantile memories requires a carefully orchestrated hippocampus-centered network process that unfolds in three stages. First, exposure to a contextual reminder of the original experience primes the hippocampal network, increasing the recruitment of neurons associated with the forgotten infantile memory (iEngram) during subsequent experiences associated with the forgotten infantile memory (“**Priming**”, **A**). Then, a reminder of the aversive stimulus experienced during infancy selectively tags iEngram neurons for offline reactivation in the following hours (“**Tagging**”, **B**). Increased iEngram activity during offline network reactivation events binds previously latent infantile memory with novel neuronal ensembles, thereby reinstating behavioral responses consistent with the original experience (“**Binding**”, **C**).

## Materials and Methods

### Animals

All animal experiments were approved by the Cantonal Veterinary Office Basel-Stadt, and all procedures were performed in compliance with its guidelines and the Swiss Veterinary Law (permit numbers 3018 and 3037). This study made use of C57BL/6JRj wild type (WT) mice, and the following transgenic strains: Fos^tm2.l(icre/ERT2)Luo^/J (TRAP2) (Cat. No. 30323, Jackson Labs), B6.129P2-Pvalb^tml(cre)Arbr^/J (PV-Cre) (Cat. No. 17320, Jackson Labs), and TRAP mice crossed with B6.Cg-Gt(ROSA)26Sor^tm9(CAG-tdTomato)Hze^/J (Flex-tdTomato) (Cat. No. 7905, Jackson Labs) or B6.Cg-Gt(ROSA)26Sor^tm40.l(CAG-aop3/EGFP)Hze^/J (Flex-ArchT) (Cat. No. 21188, Jackson Labs). All transgenic mouse lines were on a C57BL/6 background. Animals were housed in temperature- and humidity-controlled cages at 21-23 °C and 50-60% humidity, with 12 h - 12 h light-dark cycle, and with food and water provided ad libitum. All mice were group-housed with littermates and then single-housed either after surgical implantation of optic fibers or a microendoscope, or 1-3 days before the onset of behavioral experiments in adulthood. For experimental procedures, mice of both sexes were randomly and blindly allocated to experimental groups by the responsible researcher.

### Surgical procedures

#### Viral injections in pups

Two-day-old (P2) TRAP2 pups were anesthetized with 4% isoflurane in oxygen (Attane, 0.7 L/min airflow, Piramal), delivered via a custom-designed neonatal nose cone mounted on a stereotaxic frame (Kopf; see https://github.com/donatolab/isoMaskMousePups). Viral constructs were delivered via intracranial injections using a pulled glass capillary connected to a Nanoject III system (Drummond). Viral solution was injected into the hippocampus in pulses of 25 nL at 10 nL/s, with 5 s intervals between pulses. Injection coordinates were: anterior-posterior (AP): +1.0 mm; medial-lateral (ML): ±1.3 mm; dorsal-ventral (DV): −1.65 mm, relative to the junction of the medial and lateral sinuses over the cerebellum. Given the lack of calcification of the skull at that age, injections were performed without any incision of the skin or craniotomy. The following AAV constructs were used: (1) pAAV.Syn.GCaMP6f.WPRE.SV40 (Addgene #100837-AAV1; titer: 7.4 x 1012 or 1.3 x 1013 vg/mL), (2) pAAV-CAG-FLEX-tdTomato (Addgene #28306-AAV1; titer: 1.2 x 1013 vg/mL; diluted 1:50 in filtered PBS), and (3) pAAV-hSyn-DIO-hM4D(Gi)-mCherry (Addgene #44362-AAV1; titer: 2.3 x 1013 vg/mL). Viruses were mixed in a 1:1 ratio before injection, depending on the experimental condition. A total volume of 300 nL of GCaMP6f and/or FLEX-tdTomato was injected unilaterally into the right hippocampus. For experiments requiring bilateral inhibition, 200 nL of DREADD-expressing AAVs were injected into both hippocampi.

#### Viral injections in adults

Adult mice (> P60) were pre-treated with buprenorphine (Bupaq P, 0.1 mg/kg, s.c., Streuli) and atropine sulfate (0.05 mg/kg, s.c., Amino AG) 15-30 min before surgery. Anesthesia was induced with 5% isoflurane in oxygen and maintained at 1.7-2% (Attane, 0.6 L/min airflow; Piramal). A local anesthetic mix of lidocaine (< 7 mg/kg, Streuli) and bupivacaine (1-5 mg/kg, Sintetica) was administered subcutaneously over the surgical site. Mice were placed in a stereotaxic frame (1900, Kopf), and body temperature was continuously monitored and maintained using a heating pad. The scalp was sterilized with betadine (Mundipharma), and a midline incision was made. The periosteum was removed, and the skull was cleaned with betadine and sterile saline (B. Braun Surgical). Craniotomies (0.4 mm diameter drill bit, Jota AG) were drilled above the target regions. For bilateral delivery of DREADDs, PV-Cre mice received injections of pAAV-hSyn-DIO-hM3D(Gq)-mCherry (Addgene #44361-AAV1; titer: 2.2 x 1013 vg/mL) into the dorsal hippocampus (AP: −1.75 mm; ML: ±1.95 mm; DV: −1.85 mm; 350 nL per hemisphere) using a Nanoject III (Drummond). The injection rate was 10 nL/s. Craniotomies were sealed with UV-curable cement (Venus Diamond Flow, Kulzer), and the scalp was sutured (Prolene 8889H, Ethicon).

#### Microendoscopes implantation for 1-photon calcium imaging

Microendoscopes (Prism-attached GRIN lenses) were implanted 2-3 weeks before behavioral testing (as in Kveim et al.(*21*)). Mice received dexamethasone (5 mg/kg, i.m.) 4 h before surgery. After anesthesia and analgesia was induced as previously described, the skull was scored with a needle to improve adhesion, followed by application of Histoacryl (B. Braun). A 1.5 mm2 craniotomy was drilled, the dura removed, and a microendoscope (GRINtech or Inscopix; see below) was lowered adjacent to the dorsal hippocampus in the right hemisphere (prism facing posterior-medially at 45°; coordinates: AP: −1.6 mm; ML: +2.05 mm; DV: −2.3 mm, followed by −0.1 mm compression). Custom microendoscopes specifications were as follow. GRINtech microendoscope: singlet tubular GRIN lens, diameter 1.0 mm, wavelength 520nm, non-coated; prism: 1.00 x 1.00 x 1.00 mm, aluminum coating on hypotenuse. The total length of the microendoscope was approximately 4.67 mm, working distance 0.15 mm in water from prism exit surface (NA 0.4), 0.08 mm in air on the image side (NA 0.5). GRINtech GmbH, Jena. Inscopix microendoscope: cat. Nr. 1050-004601, approximate microendoscope length: 4.3 mm. The microendoscope was fixed with Venus and superglue (Loctite 415), and the surrounding skull was covered with dental cement (Paladur, Kulzer) mixed with graphite (Carl Roth, 7614). A custom metal head bar was affixed. The microendoscope was temporarily protected with Kwik-Cast (World Precision Instruments). One week later, a miniscope baseplate (Inscopix, #1050-004638) was attached under light anesthesia. The miniscope (nVue 1.0, Inscopix) was positioned until tdTomato-positive iEngram neurons were visible (red channel), and activity confirmed by green-channel calcium transients. The baseplate was secured with Venus and Loctite 415, followed by a final protective layer of Paladur with graphite. A baseplate cover (Inscopix #1050-004639) was used between sessions.

#### Optogenetic fiber implantation

Optical fibers (MFC_200/245-0.37_2mm_ZF1.25(G)_C45, Doric) were bilaterally implanted adjacent to the CA3 pyramidal layer 1-2 weeks before behavioral reinstatement. After scalp preparation as above, 0.4 mm craniotomies were made (centered at AP: −1.75 mm; ML: ±1.95 mm), the dura removed, and fibers lowered to DV: −1.85 mm. Fibers were secured with Venus and Loctite 415, followed by Paladur with graphite. A custom metal head bar was attached, and fibers were capped when not in use.

#### Postoperative care and verification

After surgery, mice recovered under a heat lamp in their home cage. Buprenorphine (0.1 mg/kg, s.c.) was administered 4-6 h postoperatively. Meloxicam (5 mg/kg, s.c., Metacam, Boehringer Ingelheim) was given once daily for 3 days. All injection sites, microendoscope placements, and fiber positions were confirmed by histological analysis.

### Behavioral protocols

#### Contextual fear conditioning

To induce the encoding of a hippocampus-dependent associative memory through contextual fear conditioning (cFC) during infancy, P19 mouse pups (“cFC P19” group) were placed in a conditioning chamber (training context (TR), Ugo Basile; tracking software: EthoVision XT, versions 14-17, Noldus), which consisted of a square box (25 x 25 cm) with a grid floor (fine spacing, electrifiable), patterned walls (stripes and squares), and a distinct odor (2% acetic acid). After 3 min of free exploration, animals received five foot shocks (unconditioned stimulus, US: 0.8 mA, 1 s each) delivered at 30 s intervals. Before and after this “cFC Training” session, pups were transported individually in a cage from their home cage to the experimental room and back. To ensure developmental comparability, animals weighing < 7 g at P19 were excluded from training. Pups were weaned at P21 and remained group-housed with littermates until single housing was required for downstream experiments. Control groups included: (1) a “Naϊve” group that remained in the home cage at P19 without exposure to cFC; (2) an adult control group (“cFC P60”) underwent cFC in adulthood (> P60) using the same training context and protocol.

To test for recent and remote memory persistence in the cFC P19 or cFC P60 groups, mice were tested for their memory expression one day and > 40d after cFC acquisition. Therefore, animals were re-introduced to the training context (CS) to freely explore for 5 minutes without shocks (no US).

#### Memory reinstatement

To reinstate a forgotten infantile memory, mice underwent a multi-day behavioral protocol adapted from (*11*), beginning at least 40 days after the initial contextual fear conditioning (cFC) in infancy. On Day 1 (“reminder CONTEXT” session, C), animals were reintroduced to the original training context (TR) for 5 minutes. No shock was delivered; the session served as a non-aversive reminder of the infantile memory. On Day 2 (“reminder AVERSIVE” session, A), approximately 24 hours later, mice were placed in a novel context consisting in a circular arena (80 cm diameter) with black walls, an electrifiable grid floor (large spacing), wiped with ethanol, and located in a different room than TR. After 4 minutes of exploration, a single foot shock (0.8 mA, 2 s) was delivered, and mice were removed 1 minute later. On Day 3 (“recall Reinstated” session, R), −48 hours after CONTEXT, mice were returned to the original training context (TR) and allowed to explore for 5 minutes without shock. To assess the specificity of the reinstated memory, a subset of animals was tested in a distinct novel context rather than the training environment. This context (termed CONTEXT B) was a small circular arena, 27 cm in diameter, with white walls and scented with 0.25% benzaldehyde diluted in 70% ethanol. To assess the strength of the memory encoded for the specific association between the circular arena and the single foot shock experienced during AVERSIVE, a subset of animals was put back into the AVERSIVE reminder arena at least 24 hours after the reminder session.

Several variations of the standard CONTEXT-AVERSIVE-REINSTATED (“C-A-R”) sequence were implemented. In a subset of animals, we tested the contribution of the CONTEXT session itself by omitting it (“no CONTEXT”), or by replacing it with exposure to CONTEXT B either only during the CONTEXT reminder session (“B-A-R”), or both during the reminder and recall session (“B-A-B”). In a different cohort, Context B was made more similar to TR by applying the TR-specific odor (“B+Odor -A-R”). These sessions, which lasted five minutes each, were designed to isolate the contextual and olfactory contributions to reinstatement. To investigate the necessity of the aversive component, a control group underwent the AVERSIVE session without receiving a shock (“no AVERSIVE”). In another variation, the foot shock was replaced with a different aversive experience: mice were allowed to freely explore an elevated transparent Plexiglas platform (12 x 12 cm, 95 cm high, wiped with ethanol) for thirty minutes. This session (“C-ELE-R”) was designed to elicit innate fear responses.

In other cohorts, the delay between CONTEXT and AVERSIVE sessions was shortened to six or fifteen hours, or the interval between the AVERSIVE and REINSTATED sessions was extended to three, seven, or twenty-eight days. To test the influence of session order, the sequence or number of events was rearranged while maintaining approximately 24-hour intervals between each session. These permutations included AVERSIVE-CONTEXT-REINSTATED (“A-C-R”), repeated AVERSIVE sessions before REINSTATED (“A-A-R”), repeated CONTEXT exposures before REINSTATED (“C-C-R”), and repeated AVERSIVE and CONTEXT sessions (“C-A-A”).

#### Extinction learning

In the “TR A-Late, 1d” and “TR A-Late, 7d” groups, animals were reintroduced to the training context at about 15 hours after the AVERSIVE reminder, and allowed to explore freely for thirty minutes without receiving a shock. Another control group (“CXT B A-Late, 1d”) explored a distinct novel context (CONTEXT B) for thirty minutes during the A-Late offline window, serving as a non-specific extinction-like experience. To assess the impact of these experiences on memory stability, REINSTATED recall testing was performed 24 hours or 7 days later.

#### Quantification of behavior

Mouse behavior was recorded with an overhead infrared camera (Basler acA1300-60gm, Basler GenICam, 25 Hz) during all behavioral experiments. Video recordings were processed offline, and memory retention and expression were quantified by the amount of time the animal spent freezing during each session. Freezing bouts were systematically identified by the EthoVision XT software, and defined as moments of complete immobility apart from breathing (pixel change < 1-2% adjusted for each mouse, for at least 2 seconds).

To assess whether freezing behavior in the “cFC P19” group reflected an uniform or heterogeneous memory response, we performed a multi-step analysis in MATLAB. First, we examined the distributional structure of the P19-trained group using a Gaussian Mixture Model (GMM). Both one- and two-component GMMs were fit with a regularization parameter of 1×10-s, and Bayesian Information Criterion (BIC) scores were computed across 10,000 bootstrap resamples. The two-component model was favored in 92% of iterations, suggesting that the data were better explained by a bimodal distribution. This conclusion was further supported by Hartigan’s Dip Test, which returned a significant dip statistic and identified a primary modal interval consistent with a concentration of high-freezing values. To define potential behavioral subgroups, we then applied k-means clustering (k = 2, 10 replicates) to the P19-trained group. This analysis revealed two subpopulations: a low-freezing group and a high-freezing group. Subsequent statistical comparisons showed that the low-freezing cluster did not significantly differ from naϊve controls (Welch’s t-test p = 0.073; Mann-Whitney U p = 0.505), while the high-freezing cluster showed robust differences (Welch’s t-test p < 0.00001; Mann-Whitney U p < 0.001). These clusters were therefore interpreted as “non-learners” and “learners” respectively. To classify additional animals from independent experimental cohorts we used a centroid-based approach, where animals were assigned to the group whose k-means cluster centroid (mean freezing value) they were closest to in Euclidean space. To incorporate the structure and variability of the original data, we also computed Mahalanobis distances between each new animal and both cluster centroids using the pooled covariance of the initial dataset.

### Labelling of engram neurons

To permanently label engram neurons, we used the TRAP (Targeted Recombination in Active Populations) approach (*18*). A stock solution of 4-hydroxytamoxifen (4-OHT, MedChemExpres, HY-16950; 100 mg/mL in DMSO, CarlRoth A994) was diluted 1:10 in sunflower seed oil (Sigma, S5007) and administered intraperitoneally (i.p.) at a dose of 50 mg/kg. To permanently label infant Engram neurons (iEngram), 4-OHT was injected immediately following cFC acquisition at postnatal day 19 (P19) in TRAP2 mice crossed with either Flex-tdTomato, Flex-ArchT, or injected with a viral vector encoding Flex-tdTomato or DIO-hM4D(Gi)-mCherry. This allowed activity-dependent labeling of iEngram neurons. To permanently label neuronal ensembles recruited during the Late offline window post-AVERSIVE reminder, 4-OHT was injected 12 h after the reminder in TRAP2 mice crossed with the Flex-ArchT line. Control groups included: (1) mice injected with 4-OHT two days after cFC (P21), without any behavioral experience around the time of injection, to label a random neuronal population active at baseline, these animals were weaned at P23; and (2) mice injected during adulthood, specifically during the C-Late and A-Mid offline windows. The TRAP system showed minimal baseline recombination (“leakiness”) in the absence of 4-OHT, consistent with previous validation of the TRAP-2 x Flex-tdTomato line (Kveim et al.,(*21*)).

### Pharmacogenetic manipulations

To activate inhibitory networks in PV-Cre/DIO-hM3D(Gq)-mCherry expressing mice, clozapine-N-oxide (CNO; Sigma, C0832) was administered intraperitoneally (1 mg/kg; dissolved in DMSO (Carl Roth, A994) and diluted to 0.1 mg/mL in saline, BBraun) 40 minutes before the behavioral session. Control animals expressing PV-Cre/DIO-hM3D(Gq)-mCherry received an equivalent volume of saline at the same time point. For pharmacogenetic inhibition of iEngram neurons in TRAP2/DIO-hM4D(Gi)-mCherry expressing mice, CNO was injected intraperitoneally during defined offline phases: 12 h after the CONTEXT session (C-Late), or 7.5 h or 12 h after the AVERSIVE session (A-Mid and A-Late, respectively). Control groups included TRAP2/DIO-hM4D(Gi)-mCherry expressing mice injected with saline instead of CNO, and animals that received CNO at A-Late but had not been virally transduced. As all control groups exhibited comparable freezing levels, they were pooled for analysis.

### Optogenetic manipulations

Mice mice were briefly secured to a head post on a planar running wheel. A splitter-branching fiber-optic patch cord (SBP(2)_200/220/900-0.37_4m_FCM-2xZF1.25(F); Doric) was connected to bilateral implanted optical fibers. Following a brief recovery period (7-10 min) in the home cage, mice were transferred to the behavioral context. Continuous light (532 nm, 10.0-12.5 mW at the fiber tip; Cobolt 06-DPL laser, controlled via Cobolt Monitor software) was delivered throughout the 5-minute session (CONTEXT, AVERSIVE, or REINSTATED sessions in different cohorts of animals). To habituate animals to the stimulation procedure, mice underwent a mock session one day prior to the reinstatement protocol, during which the patch cord was connected but no behavioral context was presented. Control groups included opsin-expressing mice that were attached to the patch cord but did not receive light, and a group in which iEngram neurons were labeled at P21 in the home cage, two days after cFC, and subjected to light delivery during the AVERSIVE session.

### Calcium imaging

Time-lapse calcium imaging of CA3 pyramidal neurons was performed in freely behaving mice using an integrated dual-color miniature fluorescence 1-photon microscope (nVue 1.0, miniscope, Inscopix). At the start of each recording day, the miniscope was affixed to the animal’s baseplate while head-fixed on a planar running wheel. Mice were then returned to their home cage for a baseline recording session that started by acquiring the tdTomato+ signal for 150 frames, followed by recording CA3 network activity in the green channel for 10 minutes, while the mice remained in their home cage. Subsequently, the animals were moved into the appropriate context for either the CONTEXT, AVERSIVE, or REINSTATED sessions without detaching the miniscope to maximize the probability of longitudinal tracking of the same neurons. Recordings were paused during handling. To capture CA3 network dynamics during memory reinstatement, calcium imaging was performed longitudinally across the CONTEXT, AVERSIVE, and REINSTATED sessions (“Online” sessions), as well as during six periods while animals remained undisturbed in their home cage (“Offline” sessions). Offline recordings were performed in a neutral room, distinct from any behavioral context, and consisted of three 30-minute sessions per day on both the CONTEXT and AVERSIVE days, corresponding to +3-5 h (Early), +7-9 h (Mid), and +13-15 h (Late) following each reminder session.

Data were acquired at 20 or 25 Hz with 1280 x 800 pixel resolution, corresponding to a 1000 x 625 µm field of view (FOV; 1 pixel = 0.765 µm2). For habituation, between 2-4 days before onset of reinstatement, the miniscope was attached once for 10 minutes and mice freely moved in their home cage. Meanwhile, values for LED power and gain were empirically adjusted for the green and red channel based on the overall brightness of the GCaMP and tdTomato signal, respectively, and held constant thereafter. LED intensity was calibrated using cumulative pixel fluorescence histograms to maintain saturation between 12-75%. The focal plane was set to maximize the number of visible neurons and was not changed across sessions.

### Processing of calcium imaging data

Calcium imaging data from the green channel were processed using the Inscopix Data Analysis Software MATLAB API (IDAS, v1.9.2, Inscopix), in combination with custom MATLAB (R2021b and R2022b, MathWorks) and Python (v3.8) scripts. As first processing step, home cage and respective behavior recording sessions acquired without miniscope detachment were concatenated and processed together. Images were spatially down-sampled twofold (final FOV: 640 x 400 pixels; 1 pixel = 1.530 µm2). Image regions devoid of neural signal were cropped, and FOVs were aligned across sessions using vascular landmarks and lens edges. High- and low-frequency spatial noise were removed using a Gaussian band-pass filter (cutoffs: 0.005-0.500 pixels). Lateral motion correction was performed frame-by-frame using the IDAS implementation. Cell segmentation was performed using constrained nonnegative matrix factorization (CNMFe) provided by IDAS. Most parameters were kept at default, with empirical adjustments per animal for minimum peak-to-noise ratio (10–20), minimum pixel correlation (0.90-0.95), and cell diameter (12-14 pixels :: 18.36-25.0 µm2). For each ROI, only pixels with fluorescence values within the top 20% of maximum intensity were included.

Calcium events were detected using a custom Python-based pipeline (adapted from Kveim et (*21*). Raw calcium traces were low-pass filtered (1st-order Butterworth, 0.5 Hz), and baseline drifts and bleaching artifacts were removed by subtracting a polynomial fit (order 1-2) to a slower low-pass filtered version (0.1 Hz cutoff). A global detection threshold was defined as the 50th percentile of the standard deviation distribution across all neurons, minimizing false positives in low-SNR traces. Events were further categorized into “up-phase” (positive slope) and “on-phase” (above-threshold segments), and retained if their durations were 2: 0.3 s and 2: 0.5 s, respectively. Signal-to-noise ratio (SNR) was calculated for each neuron as the mean amplitude of events divided by the standard deviation of the dF/Fo trace, excluding a 1-10 s window around each up-phase event. Neurons falling in the lowest 5-10% of SNR values (adjusted per recording) were excluded. In cases of oversegmentation, spatially adjacent ROIs (< 5 pixels) were compared, and the ROI with the higher SNR was retained.

Longitudinal cell registration across sessions was performed using IDAS tools, with a minimum spatial correlation threshold of 0.6. Binarized activity traces were generated by assigning a value of 1 to frames during the up-phase of calcium events and 0 otherwise. Only neurons with an average activity rate 2: 0.01 Hz in each of the three online sessions were included for further analysis.

For detection of GCaMP6+ tdTomato+ double-positive neurons, the red fluorescence channel was processed analogously. FOVs were spatially cropped and downsampled to match the green channel, motion-corrected, and averaged into a single-frame projection. To enhance signal-to-noise, a background-subtracted image was created by subtracting a 21×21 pixel filtered version from a finer 7×7 pixel filtered image. ROIs were identified by binarizing the image: pixels below the 99th percentile of intensity or with negative values were zeroed, and ROI boundaries were defined by pixels exceeding 80% of the local maximum. Overlapping ROIs in the green and red channels were matched, and those sharing 2: 20% of red ROI pixels were considered candidate double-positive neurons. All double-positive ROIs were manually verified. Identified GCaMP6+ tdTomato+ neurons were then tracked and analyzed across all online and offline sessions.

### Analysis of calcium imaging data

#### Single neurons longitudinal activation

To quantify the extent to which individual neurons modulated their activity rate in each online or offline recording sessions compared to the baseline period (*i.e.* in their Home Cage at the beginning of the day), we calculated a “Activation Score” (AS) for each neuron and session (*REFERENCE*). The AS was determined by computing the difference in activity rate (averaged across the whole session and expressed in Hz) between the specific online/offline session (SE) and the baseline activity rate in the Home Cage (HC), normalized by their sum:

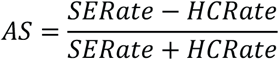

Neurons with AS 2: 0.33 (*i.e.* having double the activity rate than Home Cage) were classified as “Activated”; neurons with AS :S −0.33 (*i.e.* having half the activity rate than Home Cage) were classified as “Inhibited”; while neurons with −0.33 < AS < 0.33 were classified as “Neutral” (*21*).

#### Population co-activity across sessions

Neuronal co-activity was analyzed across nine experimental conditions (3 offline + 6 offline sessions). Each neuron was classified as active (1) or inactive (0) for each condition depending on its AS (AS 2: 0.33: active. AS < 0.33: inactive). In addition, neurons received a primary label based on their ensemble identity (CONTEXT, AVERSIVE, REINSTATED, no ensemble), and a secondary label based on their tdTomato expression (expression of tdTomato identifies iEngram neurons). Pairwise similarity between conditions was quantified using the Jaccard index, defined as the size of the intersection of active neurons divided by the size of the union. This resulted in a 9×9 Jaccard similarity matrix. A hierarchical clustering on this matrix using average linkage on the Jaccard distance (1 - Similarity) was used to assess condition groupings into clusters. For each cluster, neurons active in all member conditions were extracted. The distribution of their primary labels was compared against the global label distribution using a chi-square goodness-of-fit test. To assess the statistical significance of intra-cluster co-activity, we performed 1,000 random permutations of the neuron-condition matrix (independently shuffling neuron assignments per condition). For each permutation, the average pairwise Jaccard similarity within a cluster was recalculated. An empirical p-value was computed as the fraction of permutations that produced an equal or higher average similarity than observed.

#### Synchronous calcium events (SCEs) detection

To detect time points of population-level synchrony, we identified frames in which the number of simultaneously active neurons exceeded chance levels based on a shuffled null distribution. Binary activity traces (up-phase only) were compiled for all neurons across sessions into a matrix of size *T* x *N*, where *T* is the total number of imaging frames and *N* is the number of neurons. For each session, we computed the number of active neurons per frame in the real data. To generate the chance distribution, we independently shuffled each neuron’s activity trace across time within the session (i.e., column-wise shuffling), thereby preserving individual firing rates but removing temporal structure. This procedure was repeated 1,000 times. Across all shuffles, we aggregated the number of co-active neurons per frame to generate a null distribution of frame-wise population activity. The 95th percentile of this null distribution was used as the session-specific threshold for statistical significance. Frames in which the real number of active neurons exceeded this threshold were classified as SCEs. For each session, we quantified the frequency of SCEs and the fraction of recorded neurons participating in at least one of these events, and recorded their temporal occurrence for downstream ensemble and state-space analyses.

#### Individual neurons’ participation in SCEs

To determine whether individual neurons participated in SCEs more or less frequently than expected by chance, we implemented a cell-by-cell statistical comparison of observed versus expected activity. This analysis was based on the assumption that, under a null model of independent firing, the probability of a neuron participating in any given SCE is proportional to its overall firing rate. A matrix of binarized calcium activity was constructed for each neuron across all frames identified as SCEs. For each neuron and session, the observed participation was defined as the fraction of SCE frames during which the neuron was active. The expected participation was computed as the neuron’s overall activity rate (i.e., the fraction of all time frames in which the neuron was active) multiplied by the total number of SCE frames, assuming independence between neurons. To assess significance, the observed and expected participation values were assembled into matched matrices and compared using a z-score based approach. For each neuron *i* in condition *j*, the raw difference

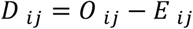

was computed, where *O* and *E* represent the observed and expected values, respectively. Differences were standardized within each condition by computing:

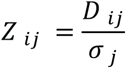

where σ*_j_* is the standard deviation of differences across neurons in condition *j*. Two-tailed p-values were derived using the standard normal distribution. Neurons with p < 0.05 were considered to participate in SCEs either above expectation (*Z > 0*) or below expectation (*Z < 0*).

#### SCEs similarity

To assess the similarity of population-wide neuronal activation patterns during SCEs, we computed pairwise cosine similarity across all SCEs within each imaging session. For each session, we extracted binary population activity vectors corresponding to SCE frames (rows) and valid neurons (columns) from the up-phase binarized calcium trace matrix. Neurons with missing values were excluded. If at least two SCEs occurred in a session, we normalized each population vector to unit length and computed a cosine similarity matrix by taking the dot product of all vector pairs. Cosine similarity values ranged from 0 (orthogonal activity patterns) to 1 (identical patterns). To summarize the overall similarity structure of SCEs, we extracted the upper triangle of the cosine similarity matrix (excluding the diagonal) and computed a histogram using 20 bins from 0 to 1. Cosine similarity matrices and their associated distributions were visualized for each session to evaluate whether population activity patterns were recurrent or variable across SCEs.

### Tissue processing and histology

For further analysis of endogenous c-Fos expression during memory reinstatement, mice were transcardially perfused with 4% PFA (Sigma Aldrich, P6148) in PBS 60 to 90 minutes after the corresponding behavior for online sessions, and +3-5h (Early), +7-9h (Mid) and +13-15h (Late) timepoints after CONTEXT or AVERSIVE reminders for offline sessions.

Before perfusion, deep anesthesia was induced with isoflurane (5% in 0.5 l/min flow) and *i.p.* injection of Ketamine/Xylazine (Ketanarkon, 250 mg/kg, Streuli; Xylazine, 2.5 mg/kg, Streuli). Brains were extracted and kept at +4 °C overnight in 4% PFA, which was then replaced by a solution of 30% sucrose (Millipore, 84100) in PBS. 40 µm coronal sections were cut using a cryostat, and collected in PBS in 24-well plates. For immunohistological staining, sections were incubated in a blocking solution consisting of 10% Bovine Serum Albumin (BSA, Sigma-Aldrich, A3912) in PBST (PBS + 0.3% Triton-X100 (CarlRoth, 3051)), for one hour at room temperature (RT). Sections were incubated in PBST with 3% BSA and primary antibodies overnight at +4 °C: Rat anti-c-Fos (1:1000, Cat. No. 226 017, Synaptic Systems, RRID: AB_2864765), Rabbit anti-c-Fos (1:1000, Cat. No. 226 003, Synaptic Systems, RRID: AB_2231974), Guinea Pig anti-NeuN (1:1000, Cat. No. ABN90P, Sigma). Sections underwent three washing steps in PBST for 15 minutes each at RT, followed by incubation with secondary antibodies (Chicken anti-rat 647 (1:500, Cat. No. A21472, Invitrogen), Donkey anti-rat 647 (1:500, Cat. No. A78947, Invitrogen), Goat anti-rabbit 647 (1:500, Cat. No. A21244, Invitrogen), Goat anti-guinea pig 488 or 568 (1:500, Cat. No. A11073/A11075, Invitrogen)) for two hours at RT in PBST containing 3% BSA. Sections were washed once with PBST and twice with PBS for 15 minutes each at RT, and mounted on microscope slides and protected by a coverslip using a custom-made anti-fade mounting medium (15% Selvol 205, Sekisui PO041611511; 30% Glycerol (AppliChem, 131339); 2.5% 1,4-Diazabicyclo[2.2.2] octan (DABCO, Sigma, D27802) in PBS). Coverslip edges were sealed with nail polish one day after mounting and stored at +4 °C. Brains with lens implants were cut to 50 µm, washed once with PBST and twice with PBS, for 15 minutes each, then mounted as above, but using custom-made anti-fade mounting medium containing 10ug/ml 4’, 6-diamidino-2-phenylindole (DAPI, Sigma, D9542).

### Confocal imaging and histology quantifications

Fluorescence images were acquired as z-stacks using an Olympus IXplore Spin Confocal Imaging Microscope System (Evident) equipped with a Yokogawa CSU-W1 spinning disk unit and a Hamamatsu ORCA-Fusion camera. The following objectives were used: a 10x air objective (UPL X APO, NA 0.4; z-stack step size: 2 µm), and a 30x silicone oil objective (UPL S APO, NA 1.05; z-stack step size: 1 µm). Laser power and exposure times were defined per fluorescent channel and kept constant throughout the study to ensure consistency across experimental conditions. Imaging parameters were initially calibrated using an adult home cage control brain and were as follows: 488 nm channel (laser power: 30%, exposure time: 150 ms), 568 nm channel (laser power: 20%, exposure time: 100 ms), and 647 nm channel (laser power: 30%, exposure time: 150 ms). To minimize variability, images from the same animal were acquired on the same day, and, when possible, samples from the same experimental cohort were imaged on consecutive days.

For spatially unbiased quantification, 3-5 coronal sections per animal were selected from the dorsal hippocampus −1.58 mm to −2.30 mm AP relative to Bregma), covering the injection or implant site. Within each section, 3-5 fields of view were acquired to include DG, CA3, and CA1 regions. Composite images were generated via stitching of adjacent fields using a custom MATLAB script (DOI: 10.5281/zenodo.7701150).

Quantification of fluorescently labeled neurons was performed in Imaris (versions 9.3.1 to 9.9.1, Oxford Instruments) using the spot detection algorithm on z-stack images. Neurons positive for c-Fos, NeuN, TRAP-tdTomato, hM4D(Gi)-mCherry, or ArchT-GFP were identified using empirically defined expected spot diameters to reflect nuclear versus cytosolic localization. An expected diameter of 6 µm was used for nuclear c-Fos+ cells, while a 7 µm diameter was used for cytoplasmic markers (NeuN+, tdTomato+, GFP+, and mCherry+). A quality threshold of 20 arbitrary units was applied to eliminate false positives, following software recommendations. Spot detection parameters were established at the beginning of the study for each marker and remained constant throughout. Only neurons with somata located in the pyramidal layer were included in downstream analyses.

Co-localization of markers (e.g., c-Fos and TRAP-tdTomato) was defined as a 3D overlap of detected spots. To distinguish biologically meaningful co-expression from chance overlap, co-labeling probability was calculated as:

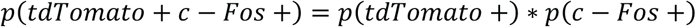

Quantifications were verified by manual inspection and normalized to the number of neurons in the region of interest. During the analysis, the researcher was blind to the experimental conditions.

### Exclusion criteria

Both male and female animals were used in all experiments. Only individuals weighing at least 7 grams at the time of contextual fear conditioning (cFC) on postnatal day 19 (P19) were included. No animals were excluded based on performance during cFC acquisition.

Following histological analysis, animals were excluded from analysis if Gi expression failed to target the hippocampal CA3 region in at least one hemisphere and instead labeled off-target regions, such as the overlying cortex. In optogenetic experiments, animals were included in the experimental group only if optical fibers were correctly placed in CA3 bilaterally.

### Statistical analysis and data visualization

Statistical analyses were performed using GraphPad Prism (versions 9.5.1, 10.0.0, and 10.2.3 for Windows and Mac, GraphPad Software, California) and MATLAB (versions R2021b, R2023b, and R2024a, MathWorks). Detailed descriptions of the statistical tests, exact p-values, sample sizes, and effect sizes are provided in Supplementary Table 2. Box plots display the 25th percentile, the median, the 75th percentile, with whiskers extending to the minimum and maximum value. The box itself represents the interquartile range (IQR), encompassing the middle 50% of the data. The line within the box indicates the median. Individual data points represent either individual animals or neurons, and are shown in each figure unless otherwise stated. Statistical significance is indicated throughout the figures and legends, as well as in Supplementary Table 2, using the following conventions: *p < 0.05, **p < 0.01, ***p < 0.001. Graphs were generated in GraphPad Prism, MATLAB, or Python, and further edited using Adobe Illustrator (versions 2022 and 2024). While no statistical methods were used to predetermine sample sizes, all sample sizes were consistent with those reported in previous studies, and post hoc effect sizes were computed and reported. Unless otherwise specified, control groups showed consistent behavior and baseline expression levels of c-Fos across experiments. Thus, control animals were pooled across cohorts and used as controls in multiple figures throughout the study.

## Bibliography

1. K. G. Akers et al., Hippocampal neurogenesis regulates forgetting during adulthood and infancy. Science 344, 598–602 (2014).

2. C. M. Alberini, A. Travaglia, Infantile Amnesia: A Critical Period of Learning to Learn and Remember. J Neurosci 37, 5783–5795 (2017).

3. J. Bevandic et al., Episodic memory development: Bridging animal and human research. Neuron 112, 1060–1080 (2024).

4. B. A. Campbell, E. H. Campbell, Retention and extinction of learned fear in infant and adult rats. J Comp Physiol Psychol 55, 1–8 (1962).

5. F. Donato et al., The Ontogeny of Hippocampus-Dependent Memories. J Neurosci 41, 920–926 (2021).

6. S. Freud, A. A. Brill, Sigmund Freud Collection (Library of Congress), *Psychopathology of everyday life*. (The Macmillan company, New York,, 1914), pp. vii, 341, 341 p.

7. A. Guskjolen et al., Recovery of “Lost” Infant Memories in Mice. Curr Biol 28, 2283–2290 e2283 (2018).

8. N. S. Newcombe, M. E. Lloyd, K. R. Ratliff, Development of episodic and autobiographical memory: a cognitive neuroscience perspective. Adv Child Dev Behav 35, 37–85 (2007).

9. A. I. Ramsaran et al., A sensitive period for the development of episodic-like memory in mice. Curr Biol 35, 2032–2048 e2033 (2025).

10. A. I. Ramsaran et al., A shift in the mechanisms controlling hippocampal engram formation during brain maturation. Science 380, 543–551 (2023).

11. A. Travaglia, R. Bisaz, E. S. Sweet, R. D. Blitzer, C. M. Alberini, Infantile amnesia reflects a developmental critical period for hippocampal learning. Nat Neurosci 19, 1225–1233 (2016).

12. T. L. Bale et al., Early life programming and neurodevelopmental disorders. Biol Psychiatry 68, 314–319 (2010).

13. B. L. Callaghan, R. Richardson, The effect of adverse rearing environments on persistent memories in young rats: removing the brakes on infant fear memories. Transl Psychiatry 2, e138 (2012).

14. C. Heim, C. B. Nemeroff, The role of childhood trauma in the neurobiology of mood and anxiety disorders: preclinical and clinical studies. Biol Psychiatry 49, 1023–1039 (2001).

15. C. Rovee-Collier, H. Hayne, Reactivation of infant memory: implications for cognitive development. Adv Child Dev Behav 20, 185–238 (1987).

16. R. Bisaz, B. Bessieres, J. M. Miranda, A. Travaglia, C. M. Alberini, Recovery of memory from infantile amnesia is developmentally constrained. Learn Mem 28, 300–306 (2021).

17. S. D. Power et al., Immune activation state modulates infant engram expression across development. Sci Adv 9, eadg9921 (2023).

18. L. A. DeNardo et al., Temporal evolution of cortical ensembles promoting remote memory retrieval. Nat Neurosci 22, 460–469 (2019).

19. P. Bekinschtein et al., Persistence of long-term memory storage requires a late protein synthesis- and BDNF-dependent phase in the hippocampus. Neuron 53, 261–277 (2007).

20. S. Karunakaran et al., PV plasticity sustained through D1/5 dopamine signaling required for long-term memory consolidation. Nat Neurosci 19, 454–464 (2016).

21. V. A. Kveim et al., Divergent recruitment of developmentally defined neuronal ensembles supports memory dynamics. Science 385, eadk0997 (2024).

22. A. Goto et al., Stepwise synaptic plasticity events drive the early phase of memory consolidation. Science 374, 857–863 (2021).

23. Y. Zaki et al., Offline ensemble co-reactivation links memories across days. Nature 637, 145–155 (2025).

24. A. Malvache, S. Reichinnek, V. Villette, C. Haimerl, R. Cossart, Awake hippocampal reactivations project onto orthogonal neuronal assemblies. Science 353, 1280–1283 (2016).

